# Optimizing data integration improves Gene Regulatory Network inference in Arabidopsis thaliana

**DOI:** 10.1101/2023.09.29.558791

**Authors:** Océane Cassan, Charles-Henri Lecellier, Antoine Martin, Laurent Bréhélin, Sophie Lèbre

## Abstract

**Motivations:** Gene Regulatory Networks (GRN) are traditionnally inferred from gene expression profiles monitoring a specific condition or treatment. In the last decade, integrative strategies have successfully emerged to guide GRN inference from gene expression with complementary prior data. However, datasets used as prior information and validation gold standards are often related and limited to a subset of genes. This lack of complete and independent evaluation calls for new criteria to robustly estimate the optimal intensity of prior data integration in the inference process.

**Results:** We address this issue for two common regression-based GRN inference models, an integrative Random Forest (weigthedRF) and a generalized linear model with stability selection estimated under a weighted LASSO penalty (weightedLASSO). These approaches are applied to data from the root response to nitrate induction in *Arabidopsis thaliana*. For each gene, we measure how the integration of transcription factor binding motifs influences model prediction. We propose a new approach, DIOgene, that uses model prediction error and a simulated null hypothesis for optimizing data integration strength in a hypothesis-driven, gene-specific manner. The resulting integration scheme reveals a strong diversity of optimal integration intensities between genes. In addition, it provides a good trade-off between prediction error minimization and validation on experimental interactions, while master regulators of nitrate induction can be accurately retrieved.

**Availability and implementation:** The R code and notebooks demonstrating the use of the proposed approaches are available in the repository https://github.com/OceaneCsn/integrative_GRN_N_induction.

## Introduction

Gene Regulatory Network (GRN) inference has the objective of deciphering the relationships between genes in the context of transcription, which can provide invaluable insight into environmental adaptation or developmental processes in living organisms. Statistical inference methods usually leverage high-throughput genomics to reconstruct those networks, in which nodes represent genes, and edges represent a relation of regulation between those genes. Because transcriptomic data are increasingly common and less costly, they are the input of choice for most statistical approaches to GRN inference.

Regression-based techniques mainly differ in their choice of regression function to link the expression of a target gene to the expression of its regulators. For example, TIGRESS [25], MERLIN [49] or The INFERELATOR [10, 38, 22] techniques implement linear parametric models for this task, while GENIE3 [27] and inspired works [21, 45, 17, 13] model non-linear relations via Random Forests (RFs) or, more broadly, ensembles of trees. Once regression models are fit, they allow the extraction of the influence of each regulator over each gene, and the strongest pairs are assembled to form a final sparse GRN.

Given the under-determined nature of GRN inference from expression alone, using additional sources of data can guide the choice between several regulators explaining expression data equally well. Complementary omics have already been used in addition to gene expression to enhance GRN inference, such as TF binding experiments (mostly ChIP-Seq) or Transcription Factor Binding Motifs (TFBM) [32, 39, 1, 18, 46, 17, 22], knock-outs and protein-protein interactions [45] or chromatin accessibility [18, 42].

In a linear context, prior information can be integrated to GRN inference by modulating the penalty strength for each TF during the estimation of regularized models with a weighted version [9] of the LASSO [57] and many variations (*e*.*g*. ElasticNet [63]) [16, 24, 56, 42, 22] or by making use of a Bayesian prior [24, 53, 22]. An in-depth approach explored a resolutive range of data integration strengths with a weighted ElasticNet, and then choose a value maximizing effective data integration [24]. More recently, the StARS approach proposing to increase the stability of non-oriented GRN inference in Gaussian Graphical Models [36] was adapted for the LASSO in order to select a small subset of robust regulators for each gene in oriented GRN inference [42]. In that work, the integrated priors are TFBMs in accessible chromatin, and three values of prior reinforcement modulating penalty strength are investigated. The moderate one is chosen to maximise the area under the precision and recall curve against a CHIP-Seq gold standard.

Regarding non-linear regression, iRafNet [45] proposed a Random Forest (RF) based procedure. It consists in weighting the random sub-sampling of regulators during trees elongation so that the chance of regulators supported by prior knowledge to get chosen at decision nodes is increased. This has the effect of inflating the importance metric of interactions supported by the chosen prior [45]. The weights controlling the contribution of prior data to expression is provided by a predefined function, specific to each type of prior, but without specific tuning. This strategy was further adapted to time series expression data in the OutPredict method [17] as an extension of both a dynamic version of GENIE3 [21]and iRafNet.

Existing integrative regression models have a great potential to predict GRN from several types of omics. However, fine tuning the contribution of prior data to expression is rarely explored. When it is, the choice of prior integration strength is set using of a gold standard that is either identical to the integrated prior [24], or of a related nature [42] (in that latter paper, the prior information of binding motifs is necessarily correlated to ChIP-seq validation data).

In this work, we propose DIOgene (Data Integration Optimization for gene networks) for a more robust and independent calibration of prior data integration strength, based on effective data integration, gene expression prediction accuracy, and a simulated null hypothesis. Moreover, to reflect the specificities of gene regulation, we propose to tune data integration strength specifically for each gene, in contrast to previous works that traditionally enforce the same integration intensity for all genes [53, 24, 45, 42, 17, 22].

In order to represent the most common methods in the field of integrative GRN inference both in the linear and non-linear cases, we illustrate our results using a weightedLASSO and weightedRF model. The proposed approaches are applied to the modelling of the transcriptomic response to nitrate induction in the roots of *Arabidopsis thaliana* [59] using TFBM in gene promoters. as prior information. With this study, we hope to open a reflection about data integration and evaluation practices in the field.

## Material and Methods

We present in the following sections weightedLASSO and weightedRF, two integrative GRN inference procedures based on TFBM prior information (Figure 1). The TFBM prior matrix Π gives, for each regulator-target pair (*r, t*) a prior value Π_*r,t*_ ∈ [0, 1] defined as:

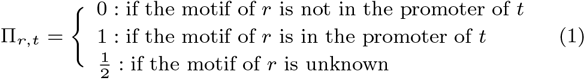

Throughout this study, the parameter *α* will be used to tune data integration strength in GRN inference : its value ∈ [0, 1] controls the contribution of TFBM information to expression data. When *α* = 0, expression alone is used, while at *α* = 1, gene expression is used to chose only between regulators possessing a TFBM in the target gene. The integration of *α* in the two inference algorithms is detailed below, as well as our original permutation based-procedure to determine the optimal *α*, in a gene-specific manner. The Implementation of this integration tuning in weightedLASSO and weightedRF is detailed below.

**Fig. 1:**
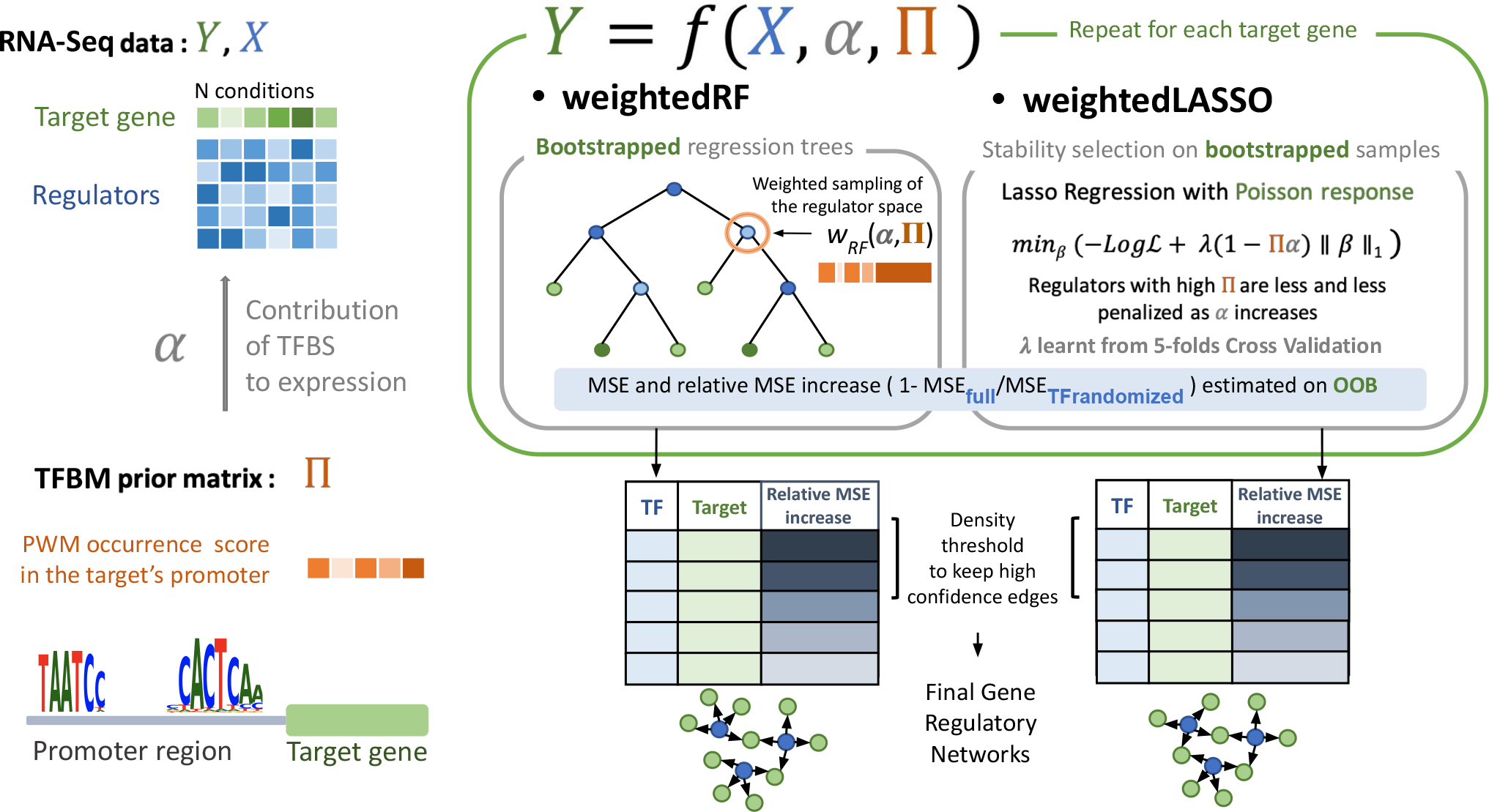
Overview of weightedRF and weightedLASSO integrative GRN inference procedures. Input data for a given target gene is the vector of expression of this gene *Y*, an expression matrix for the regulators *X*, and a TFBM scoring matrix Π. Π contains information about the presence or absence of the regulators PWM in target gene’s promoter. *α* is a parameter controlling the force of TFBM integration to expression. For each target gene, a regression model is fit to *Y* with weightedRF or weightedLASSO using X as predictive variables, and favoring the attribution of high influences to regulators with high values in Π. For weightedRF, this prioritization is executed by a weighted subsampling of the regulators space when elongating the regression trees, while it is achieved by differential shrinkage combined to stability selection is weightedLASSO. Once all regulator-gene pairs have been ranked based on their influence in the regression models, final GRN are built by selecting the number of strongest interactions providing a desired network density.

### Generalized Linear Model with Weighted LASSO (weightedLASSO)

As RNA-Seq experiments generate count data, we model the expression of a target gene *t* in the condition *i* as a Poisson-distributed variable *Y*_*t,i*_ ∼ 𝒫(*μ*_*t,i*_). The parameter of the Poisson distribution *μ*_*t,i*_ is estimated on the log scale as a linear combination of the expression values of the regulator genes, with *x*_*r,i*_ the expression level of regulator *r* in condition *i*:

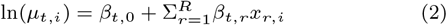

We employ a LASSO penalty in order to overcome the high-dimensional setting and to select the most predictive regulators. In addition, we propose to use differential shrinkage in order to favor the selection of TFBM-supported variables. Differential shrinkage for the LASSO (weighted LASSO) allows to modulate the penalty strength of each variable individually, in a way that regulators with a binding motif in the target’s promoter are less penalized during model adjustment. We model differential shrinkage using specific penalty coefficients *w*_*t,r*_ ∈ [0, 1] defined as a linear function of the TFBM prior Π_*t,r*_ and *α* (Figure 2a):

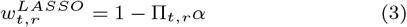

For each target gene *t*, the function to minimize for model estimation is thus:

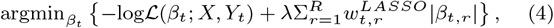

where
*λ* controls the overall strength of the penalty,

**Fig. 2:**
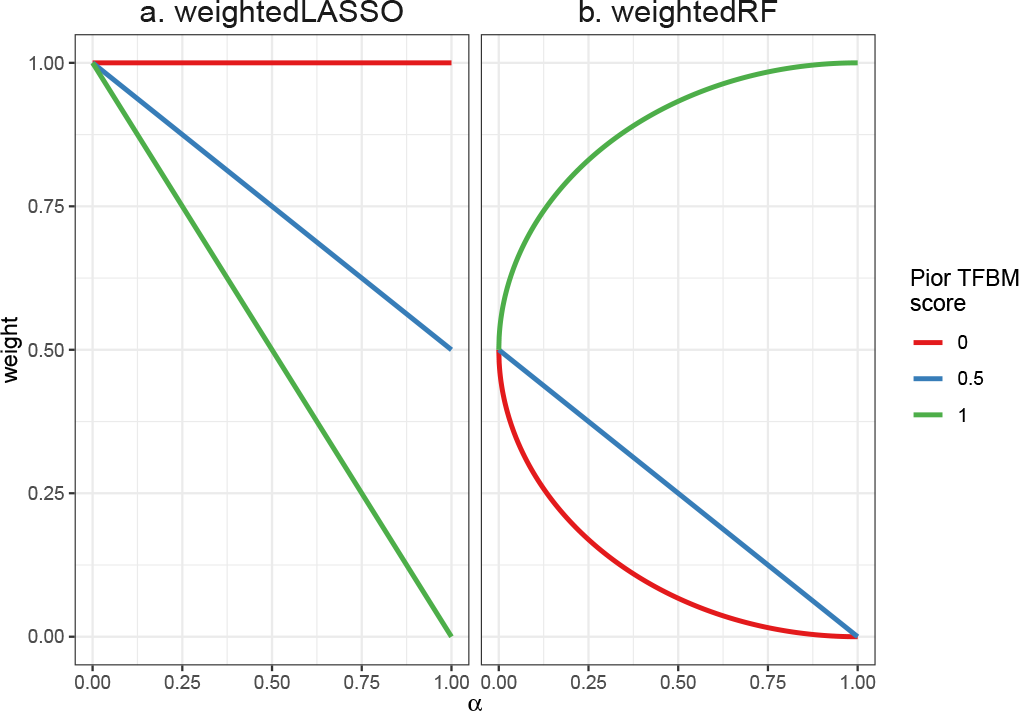
Functions linking prior TFBM scores (Π_*rt*_) to regulator-specific weights during model estimation, depending on integration strength *α*. **a**. In weightedLASSO, the penalty strength of the LASSO decreases with *α* when Π_*rt*_ *>* 0 (Equation 3). **b**. In weightedRF, the sampling weight at regression tree nodes increases with *α* when Π_*rt*_ = 1, and decreases otherwise (Equation 5). Weights are normalized between 0 and 1. In practice, these functions allow that in both cases, when *α* = 1, variable selection is performed among regulators with Π_*rt*_ = 1 only (Figure S1).

*R* is the total number of regulator genes,

*N* is the total number of expression measurements for *t*

log ℒ(*β*_*t*_; *X, Y*_*t*_) is the log-likelihood function.

The value of *λ* is learned from 5-fold cross validation. We relied on the `glmnet` [19] implementation of the LASSO, with the `penalty.factor` argument specifying differential shrinkage weights. In practice, for genes with a number *P* of TFBM-supported regulators exceeding the total number of experiments considered *N*, the model cannot be estimated at *α* = 1. We thus set *α* to 1 − *ϵ* instead of 1, with *ϵ* = 10^−4^.

To further reduce over-fitting problems and improve robustness, we also included a bootstrap procedure to the LASSO [6, 41, 37], as already used in some GRN inference approaches using a linear model [25, 42]. Hence, instead of fitting a single generalized linear model, *S* models are adjusted on a bootstrapped version of the data as follows:

1. *N* observations are sampled with replacement from the *N* available experimental conditions (bootstrapping).
2. The *N* bootstrapped observations are randomly partitioned into 5-cross validation folds. We ensure that duplicated observations during bootstrapping are grouped within the same fold.
3. A model is fitted by minimizing Equation (4) during cross-validation, allowing to learn the value of the sparsity, *λ*_1*se*_. *λ*_1*se*_ is the largest value of *λ* in the *λ* grid less than one standard deviation away from the value of *λ* minimising prediction error on the cross-validation test folds.

In this study, results are presented for *S* = 50.

### Weighted Random Forests (weightedRF)

Non-linear regressors such as ensembles of regression trees model combinatorics of regulators and complex relations between a target gene and the expression of its regulators. As inspired by iRafNet [45], we model data integration in RF by increasing the use of regulators supported by a binding motif in the decision nodes of the regression trees. A weightedRF is inferred for each target gene *t*. At each decision node, the most discriminating regulator is chosen among a subset of 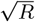 regulators. This subset, traditionally equiprobably sampled from all the regulators, is submitted here to a weighted sampling where the weights encode prior knowledge about the regulators, growing with data integration strength *α* and prior value Π_*r,t*_. Regulators with a high prior value are more likely to get chosen among the 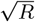 regulators tested to create a decision node. More formally, we define that the chance of a regulator *r* to get picked in the decision node for the target gene *t* is proportional to the weight 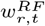(Figure 2b):

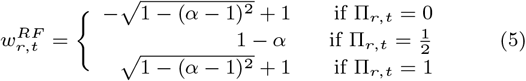

In practice, these functions allow that in both cases, when *α* = 1, variable selection is performed among regulators with Π_*rt*_ = 1 only (Figure S1). One weighted tree is learned for each of S bootstrapped samples. In this study, results are presented for *S* = 2000. The *S* trees are then aggregated into one RF per target gene. We implemented the new weight function upon the iRafNet R and C++ code, along with the possibility to restrict variables to regulators in regressions, which was not permitted initially, and with the addition of a new feature importance metric (Equation 8).

### Error in predicting gene expression (MSE)

We assess the ability of weightedRF and weightedLASSO to accurately predict gene expression via the Mean Squared Error (MSE) metric. It is measured on Out Of Bag conditions (OOB) that were left out of the bootstrapped samples (and consequently not used in weightedRF and weightedLASSO models training). For weightedLASSO, the MSE for a target gene *t*, and a given *α* is

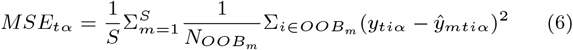

*m* corresponds to one of the *S* bootstrapped LASSO models,

*OOB*_*m*_ refers to all conditions *i* that are OOB for *m*,

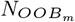 refers to the cardinal of *OOB*_*m*_,

*ŷ*_*mtiα*_ is the prediction of *m* on the condition 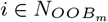.

For weightedRF, the MSE for a target gene *t* and *α* is

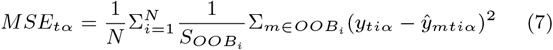

where
*i* corresponds to one of the *N* experimental conditions,

*OOB*_*i*_ refers to all trees *m* for which *i* is OOB,

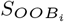 refers to the cardinal of *OOB*_*i*_,

*ŷ*_*mtiα*_ is the prediction of 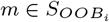 on the condition *i*.

### Importance of a regulator

Previous RF-based models [27], including iRafNet [45] which inspired weightedRF, used the traditional node purity metric. However, it is tailored for tree-based approaches only, and was shown to be not suitable to interpret variable importance in RFs in the presence of dependencies and interactions [52, 43]. We therefore define a common importance metric for both for weightedLASSO and weightedRF. This importance measure between a regulator *r* and its target *t* is given by the relative increase of MSE measured on the OOB (Equation 6 and 7), induced by shuffling the expression values of *r* when making the prediction *ŷ*_*mtiα*_ :

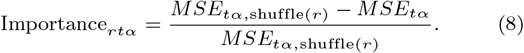

We normalize the MSE difference by *MSE*_*tα*,shuffle(*r*)_ to ensure that this statistic is comparable between different target genes and is included in the [0, 1] interval as similarly proposed in previous studies [24, 22].

### Effective Data Integration (EDI)

In order to measure the direct consequence of modulating data integration through *α*, we introduce the notion of Effective Data Integration (EDI), that reflects the importance of TFBM-supported regulators in the predictions of a regression model. For a target gene *t*, regulators are ranked by increasing values of importance, and the EDI is the average position in this ranking of TFBM-supported regulators, *i*.*e* the regulators for which Π_*r,t*_ = 1.

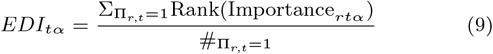

EDI is close to 1 (resp. R, the total number of regulators) when all regulators with a motif have low (resp. high) importance. We expect that increasing *α* will increase the importance values of TFBM-supported regulators, and thus increase EDI.

### Gene-specific optimisation of *α* (DIOgene)

Recall that both in linear and non linear models, *α* is the parameter controlling the extent to which TFBM contribute to model estimation relatively to gene expression, with possible values between 0 and 1. Choosing the value of *α* and consequently acting upon EDI is instrumental: it reflects strong modelling assumptions and has tangible impacts on inferred GRN. Given that enforcing data integration interferes with model estimation based solely on the minimization of the error, we can expect that this may also deteriorate the predictions accuracy. The foundation of DIOgene is that we should integrate the prior information of TFBM only if the prediction performance is not deteriorated too much.

In order to define what is an acceptable loss of prediction performance, we create a synthetic null hypothesis that will provide a reference for comparison. In this simulated null dataset, we break the link between gene expression and TFBM scores by randomly unmatching the expression profiles between regulators. A regulator then keeps its correct TFBM, but is attributed the wrong expression profile. In such a synthetic baseline, there is theoretically no joint information to be learned from the combination of expression and TFBM, and increasing data integration strength can only provide uninformative TF-gene interactions.

We can then assess prediction error (Equations 6, 7) on true data relatively to this synthetic dataset at comparable EDI (Equation 9) values for various value of integration strength *α*. In order to identify the appropriate amount of TFBM knowledge to inform GRN inference, we propose that the optimal value of *α* for target *t* (hereafter denoted as *α*_*t,opt*_), is chosen where true prediction error is most reduced as compared to the error committed under the simulated null hypothesis. This corresponds to a level of data integration where TFBM incorporation in the model provides a sufficient improvement of prediction over the shuffled baseline.

In order to build these curves, we run weightedRF and weightedLASSO for values of *α* ranging from 0 to 1 with a step of 0.1, and collect the variations of EDI and MSE depending on *α*. In order to capture the whole range of possible variations in the simulated null data but also the stochasticity inherent to the models, we run a large number of repetitions of weightedRF and weightedLASSO (resp. 100 and 50 repetitions) on both the true and simulated datasets. The curves *MSE*_*t*_(*α*) and *EDI*_*t*_(*α*) can then be assembled into one *MSE*_*t*_ = *f*_*t*_(*EDI*_*t*_) curve to optimize *α* (see several gene examples in Figure 3 for weightedRF, and Figure S2 for weightedLASSO).

**Fig. 3:**
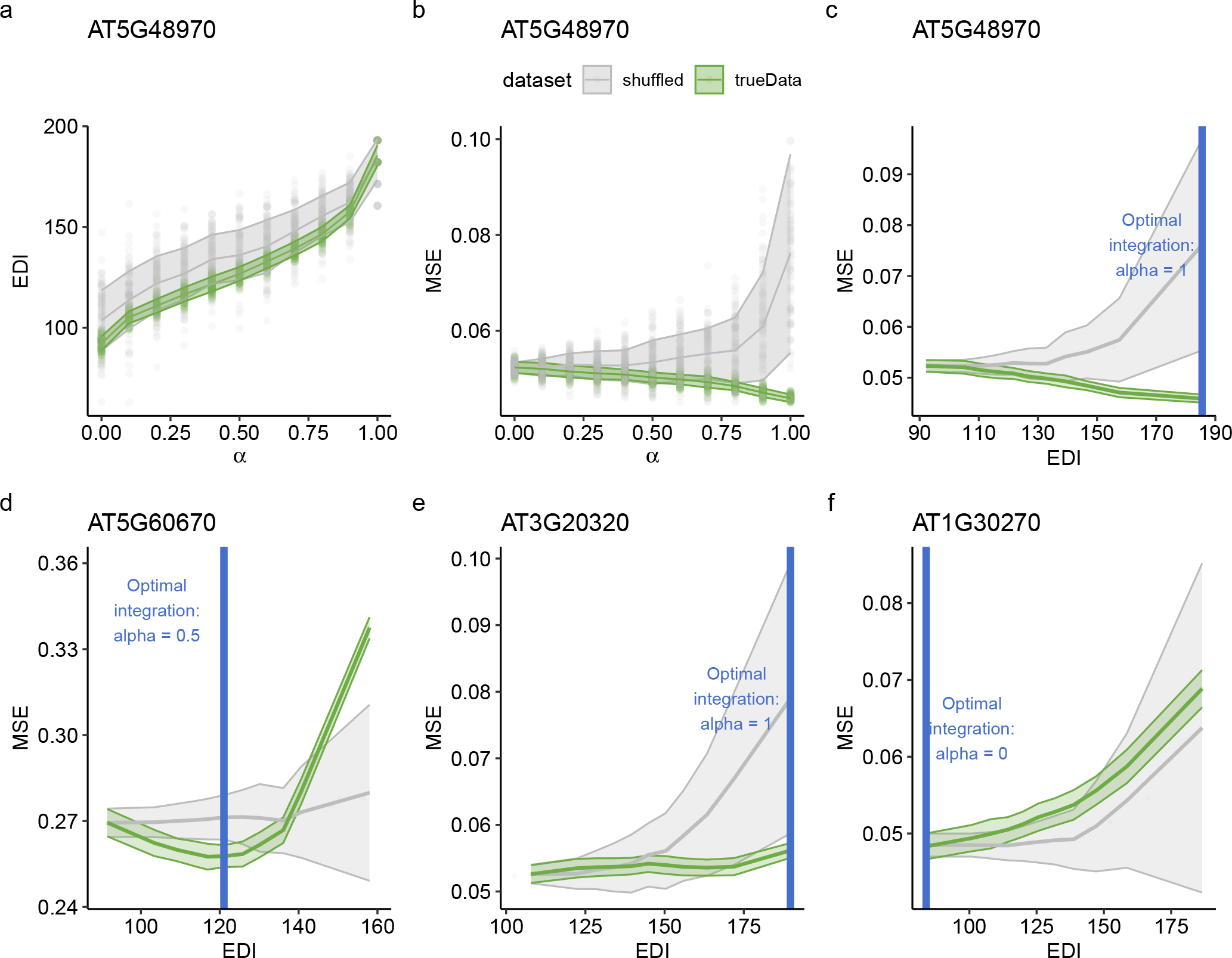
Gene-specific data integration with DIOgene is tuned by monitoring model performance variation relatively to a synthetic null hypothesis. For the target gene AT5G48970 : **a**. the EDI depending on *α*, **b** the MSE depending on *α*, **c**. the MSE depending on EDI (from panels a and b). The proposed gene-specific *α*_t,opt_ is the value for which the MSE is most reduced as compared to the shuffled baseline, represented as a vertical blue line (Equations 10 and 11). **d**,**e**,**f** : the MSE depending on EDI for three other gene examples, representing different scenarios of data integration and thus different values of *α*_t,opt_. The trends are shown for weightedRF on true data (green) and shuffled datasets where TF expression profiles were randomly unmatched from their motif (grey). For each value of *α*, 100 models were run and the standard deviation around the mean is represented. The MSE is normalized by the variance of the target gene expression. Similar scenarios emerge in the linear model weightedLASSO (Figure S2).

Formally, the normalized difference in MSE between true and shuffled datasets for a value of *α* is measured by Δ_*tα*_

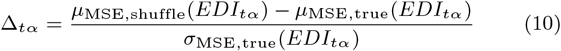

with

*μ*_MSE,true_(*EDI*_*tα*_) the mean MSE at *EDI*_*tα*_,

*σ*_MSE,true_(*EDI*_*tα*_) the standard deviation of MSE at *EDI*_*tα*_,

*μ*_MSE,shuffle_(*EDI*_*tα*_) the mean MSE on the null dataset interpolated at *EDI*_*tα*_,

*α*_*t*,opt_ is then the value of *α* that maximizes Δ_*tα*_ on a certain condition:

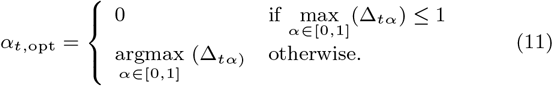

When 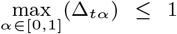, no level of data integration seems appropriate with regard to the shuffled null data, hence *α*_*t*,opt_ is set to 0. As an illustration, the vertical blue line in Figure 3 (c-f) indicates *α*_*t*,opt_ in 4 gene examples.

### Edges selection

For each target gene t, a regression model is learned with *α*_t,opt_ (Equation 11). Then, a final sparse GRN is built by selecting the edges associated with the strongest regulator-target interactions as given by the importance metric (Equation 8). Density is a common topological descriptor of biological networks: the sparsity of GRN confers them low density values, typically between 0.001 and 0.1 [33, 31, 12, 26]. A classical strategy in GRN inference is to select edges satisfying a biologically relevant user-specified network density defined as 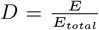, with *E* being the number of edges in the inferred network, and *E*_*total*_ = *R*(*T* − 1) being the total number of edges in a complete oriented GRN containing *R* regulators and *T* genes [13]. The number of top-ranked edges to select in order to satisfy a density *D* is thus

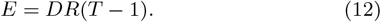

### Evaluation against experimental interactions

Let’s define 𝒢 as the set of experimental regulatory interactions (gold standard) restricted to interactions involving genes given as input for GRN inference. ℰ is the set of inferred oriented interactions restricted to TFs studied in the gold standard. The other inferred interactions can neither be confirmed nor falsified, and are thus not taken into account here. Two standard metrics for assessing the quality of a GRN are:

1. **Precision**, the fraction of edges in ℰ present in 𝒢:

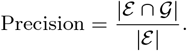
2. **Recall**, the fraction of edges in 𝒢 retrieved by GRN inference:

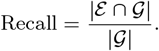

### Experimental data analysis in Arabidopsis

#### Gene expression dataset

As a case study for GRN inference, we chose the transcriptomic root response to nitrate induction in the model plant *Arabidopsis thaliana* [59]. This dynamic response has the advantage of being already well characterized, and used in other previous developments to chart regulatory networks [59, 11, 17]. Continuing efforts to uncover these regulatory mechanisms is of great agricultural interest, as nitrate is the main source of nitrogen used by most plants. Gene expression was measured in seedling roots at times 0, 5, 20, 30, 45, 60, 90, and 120 minutes after nitrate or control treatments^1^. Each combination of time point and treatment was measured in three replicates, resulting in a total of *N* = 45 samples. The RNA-Seq raw counts were normalized via the TMM method [48], and lowly expressed genes were removed prior to differential expression analysis. We selected differentially expressed genes responding to nitrate induction in time by testing the interaction terms between nitrate treatment and time modelled as natural splines, as proposed in the original article that produced this dataset [47, 59]. A total of *T* = 1, 426 genes had FDR adjusted p-values under 0.05. Among those 1,426 genes, *R* = 201 are annotated as transcriptional regulators. These nitrate-responsive genes and regulators are taken as input for GRN inference (Tables S1 and S2).

#### TFBM dataset

We define a promoter sequence in Arabidopsis as the sequence spanning -1,000 bp and +200 bp around gene start as defined in TAIR10, because this interval has been estimated to contain 86% of binding sites in plants [62]. TFBM information, encoded by Position Weight Matrices (PWM), was retrieved from the JASPAR database [14] and The Plant Cistrome Database [44].

Among the 201 nitrate-responsive regulators, 70 regulators were associated with a known PWM in the union of these databases. FIMO [23] was used to find occurences of these 70 PWMs with a significance threshold of 1e^−4^ in promoter sequences. When several occurences were identified in one promoter, the maximum score was kept.

The TFBM occurrences within nitrate-responsive gene promoters form a prior TFBM network between TFs and targets in which a target promoter harbors in average 23 TFBMs (Figure S4a) and a given TFBM is found in approximately 500 target promoters in average (Figure S4b).

## Results

### Optimal TFBM integration strength differs strongly between target genes

We ran weightedRF and weightedLASSO for values of *α* ranging from 0 to 1. We first observe that both weightedRF and weightedLASSO effectively incorporate TFBM information during their estimation, attributing higher importance measures to TFBM-supported variables as *α* increases. This is supported by EDI curves smoothly increasing with *α* (Figure S5). When applying a density threshold to build sparse GRN, we also observe that increasing *α* leads to the selection of edges with more and more TFBM support. At *α* = 1, TFBM support equals 1 meaning that, at the maximal level of data integration, GRN are restricted only to interactions supported by a TFBM (Figure S1).

An overview of the MSE profiles depending on *α* for all nitrate-responsive genes reveals a lot of diversity in how model performance can be driven by data integration strength, foreshadowing the usefulness of a gene-level procedure (Figure 4a). We thus applied DIOgene to optimize TFBM integration at the gene level (Equations 10 and 11). This confirmed that depending on the target genes, enforcing data integration has different effects on the predictive capabilities of the regression models, both in absolute error and relatively to the simulated null hypothesis. Very interestingly, for several genes like AT5G48970, increasing the EDI leads to a reduced MSE on test samples (Figure 3c). This illustrates that data integration can effectively guide the choice of variables toward more robust and meaningful regulators, allowing the model to better predict gene expression in unseen conditions. In this case, data integration can often be pushed to its maximal intensity, given that the maximal divergence from the simulated null data occur at *α*_t,opt_ = 1. For several other genes, for example AT5G60670 (Figure 3d), the strongest improvement over the shuffled baseline is achieved for an intermediate value of *α* (0.5 in Figure 3d). For genes like AT3G20320, there is no reduction of MSE due to data integration, however DIOgene sets *α*_t,opt_ to 1 because the MSE reduction in comparison to the shuffled baseline is sufficient (Figure 3e). Finally, the MSE of target genes can be increased by TFBM incorporation while showing no improvement over the simulated null data, like for instance AT1G30270, where *α*_t,opt_ is set to 0 by our procedure (Figure 3f).

**Fig. 4:**
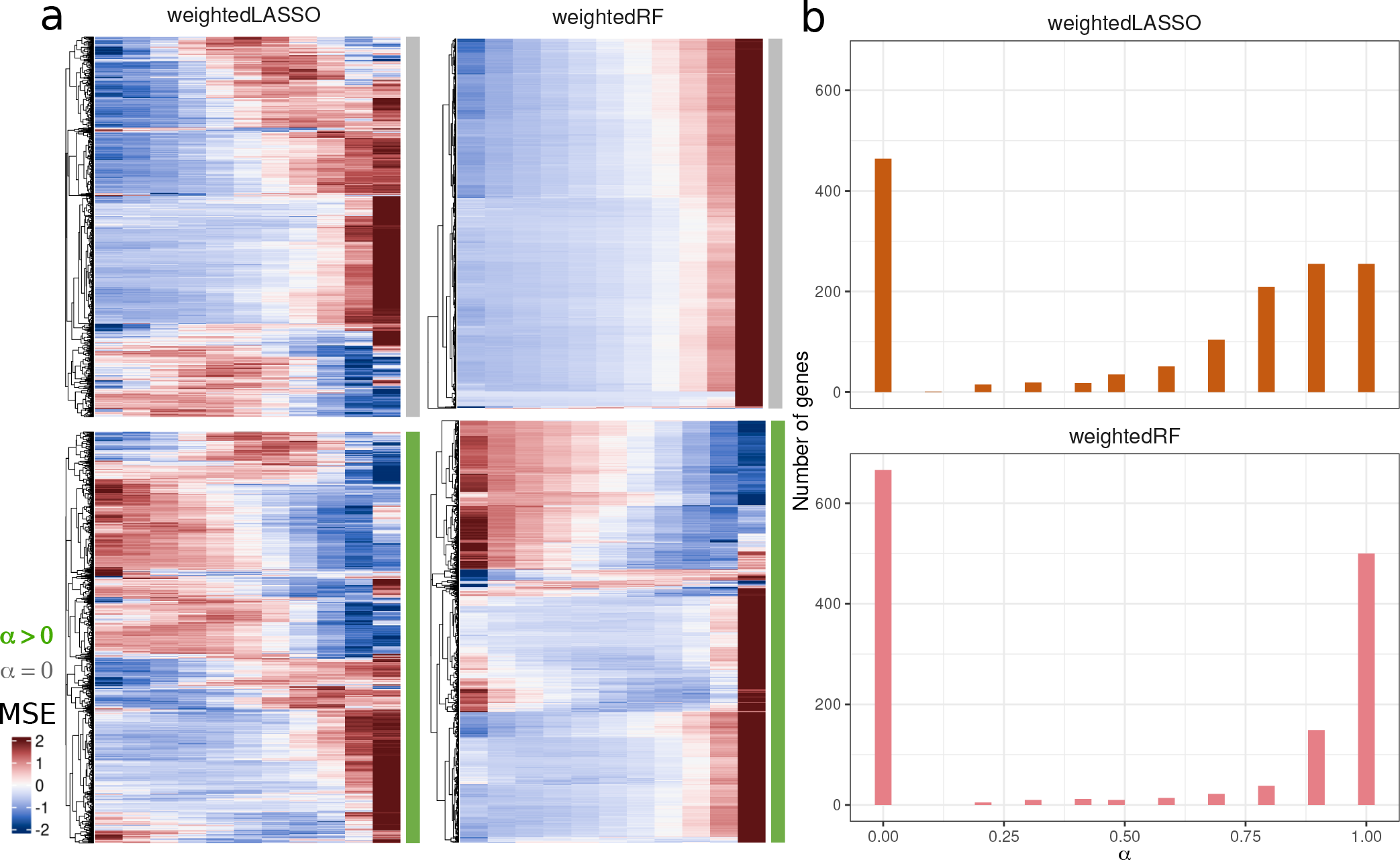
Gene-specific tuning of TFBM integration in the 1,426 nitrate-responsive genes with DIOgene leads to diverse MSE behaviors and integration intensities. a. Scaled MSE (z-score) in weightedLASSO and weightedRF on true data depending on *α* for two types of genes: genes with *α*_opt_ = 0 (grey) and genes with *α*_opt_ *>* 0 (green). b. Distribution of *α*_opt_ values for the 1,426 nitrate-responsive genes in weightedLASSO and weightedRF.

The application of DIOgene to all nitrate-responsive genes led to one *α*_t,opt_ value per target gene. Among the 1,426 input genes, the number of genes for which TFBM information is integrated to expression, *i*.*e α*_t,opt_ *>* 0, was 962 for weightedLASSO, and 760 for weightedRF. Overall, 572 genes have *α*_t,opt_ *>* 0 in both weightedLASSO and weightedRF. This significant intersection (Figure S6) indicates that the two models mostly agree on a group of genes for which data integration is beneficial, even though specificities remain. The distribution of *α*_t,opt_ for the 1,426 nitrate-responsive genes reveals that, similarly for the two models, a large pool of genes do not benefit from data integration according to our criterion (464 and 666 for weightedLASSO and weightedRF, respectively, Figure 4b). This suggests that data integration can often lead to a significant deterioration of model predictive capabilities as compared to our permuted control : in this case, our approach will leverage gene expression alone. These genes are further studied in the Discussion.

### Properties of GRN inferred with gene-specific optimisation of data integration

Moving from the individual behavior of genes in response to data integration, we now measure global properties of GRN inferred using DIOgene. In order to evaluate the added value of tuning TFBM contributions in a gene-specific manner, we compare DIOgene GRN to GRN inferred with a parameter *α* identical for all genes, as done in previous approaches [24, 45, 42, 22]. All GRN were built with a target density of 0.005, resulting in a total of 1,432 edges (Equation 12).

### DIOgene provides a trade-off between accurate gene expression prediction and agreement with a DAP-Seq experiments

We rely on three metrics to assess inferred GRN qualities: the median MSE across the 1,426 nitrate-responsive genes together with GRN precision and recall against an experimental gold standard of *in-vitro* binding interactions (DAP-Seq) [44].

First, the median MSE of GRN optimized with a global *α* displays a marked increase as the contribution of TFBM is reinforced (Figure 5a). This is in agreement with the previous observation that, for a majority of genes, TFBM deteriorate model predictions (Figure 4). Second, reinforcing TFBM contributions in GRN models equally for all genes increases both precision and recall against DAP-Seq interactions (Figures 5b, S7 and S8). Noteworthily, both models display a strong increase in precision with *α*, especially between *α* = 0 and *α* = 0.1, and weightedRF demonstrates a clear advantage over weightedLASSO, with a precision as high as 0.45. Both models outperform the precision of the prior PWM network for *α >* 0, indicating that using expression data to choose relevant links from all TFBM-supported interactions helps predicting actual TF binding, even in an *in-vitro* context. Recall values between the two models exhibit no clear differences. Thus, in globally optimised GRN, increasing data integration strength improves precision and recall, but necessarily comes with a deterioration of model predictions of the target gene expression (Figure 5a).

**Fig. 5:**
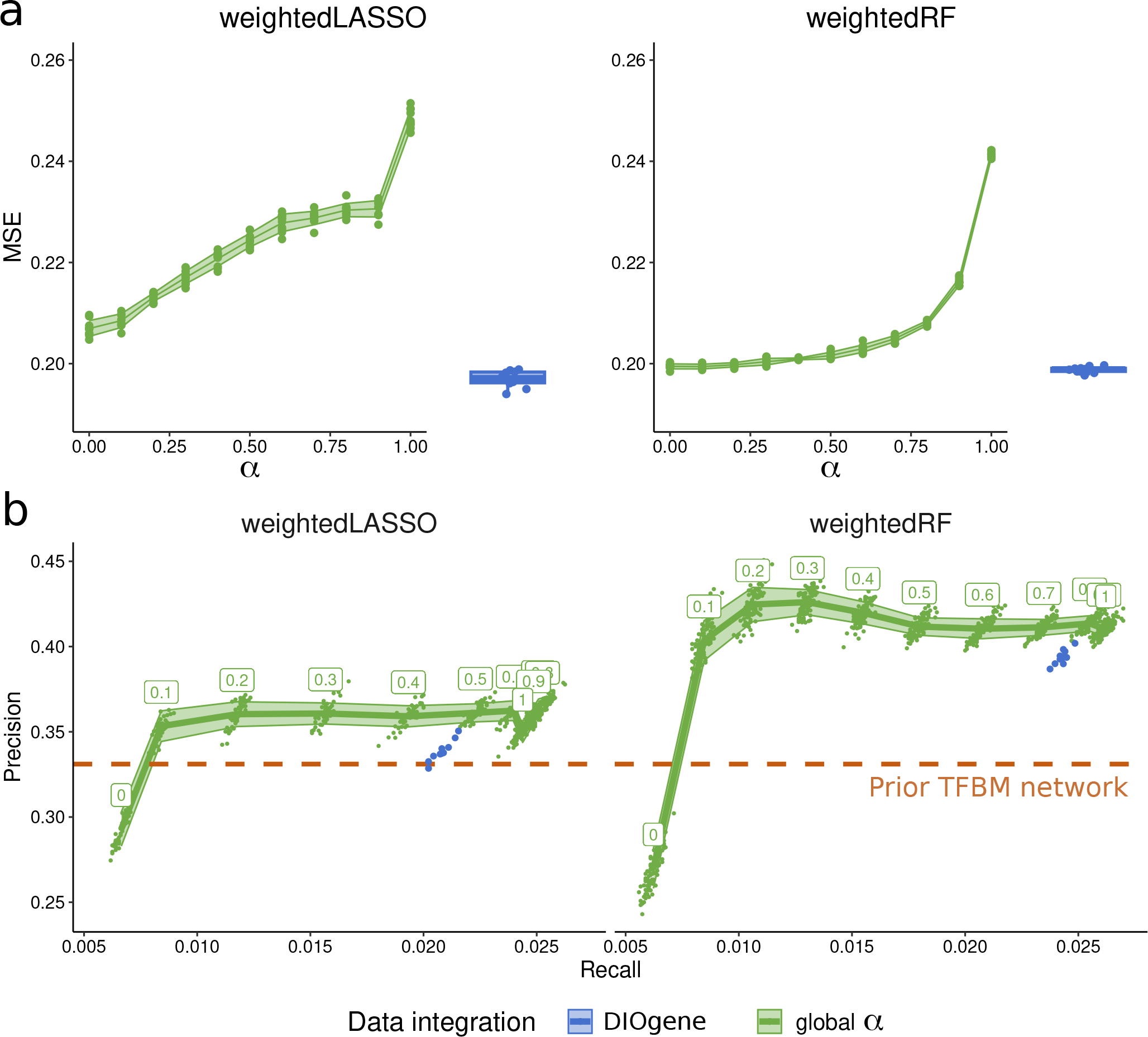
Gene-specific integration of TFBM to gene expression with DIOgene optimises model performance and secures a satisfying intersection with binding experiments. For increasing values of *α* (green) and for the proposed gene-specific optimisation of *α* (blue), we show the median MSE (normalized by the variance of the target gene) of the nitrate-responsive genes (a) and precision as a function of recall in the inferred GRN (1,432 edges, density = 0.005) against DAP-Seq interactions (b). The precision of the prior TFBM network of nitrate-responsive genes (31956 edges, density = 0.32) is overlaid in orange. Its recall is not shown because fairly comparing the recall of GRN requires that they have the same number of edges.

In contrast, gene-specific optimization of *α* with DIOgene offers a trade-off between these indicators. In fact, for both models, it provides a median MSE lower than any median MSE obtained with a global *α* (Figure 5a, blue). At the same time, GRN obtained with DIOgene achieve near-optimal precision and recall, as compared to global *α* curves. In this context, we actually argue that it is desirable to tolerate sub-obtimal precision and recall results while prioritizing low MSE, for three reasons: First, the MSE is specific to the conditions and cell lines used for transcriptome collection, which is not the case of DAP-Seq, that is agnostic of environmental and cellular context. Second, it is important to note that precision and recall also increase with *α* in shuffled datasets, where the wrong expression profile are randomly attributed to TFs (Figures S7 and S8). This illustrates that these statistics can be increased simply by enforcing data integration, even when gene expression data is uninformative. Finally, DAP-Seq and other external gold standards are often scarce and costly. Unlike the MSE, that can be measured for all genes, gold standards usually contain only a fraction of all TFs present in an inferred GRN, making it impossible to evaluate a large part of predicted interactions. We thus think that, in the context of this study and similar ones, precision and recall are unfit to properly tune, alone, the amount of a complementary omic source to incorporate into GRN inference.

### DIOgene outperforms the minimal MSE approach

Because our TFBM integration approach is based on the MSE metric, we evaluated the benefit of optimising the MSE divergence from a shuffled baseline, over a simpler approach that would minimize the MSE directly.

First, the two approaches agree on setting *α >* 0 for a large group of genes (790 and 611 for weightedLASSO and weightedRF, respectively). They also both set *α* = 0 for 224 and 538 genes. Thus, our scheme and the minimal MSE approach perform data integration on globally similar sets of genes.

In contrast, some genes are set to *α* = 0 by our approach but not by the minimal MSE (240 and 128): these genes reach a minimal MSE for *α >* 0, but do not diverge sufficiently from the synthetic null hypothesis, and are thus removed from the data integration set by our approach (see two examples in Figure S9).

Other genes are set to *α >* 0 by our approach and to *α* = 0 by the minimal MSE (172 and 149). These genes typically display an increasing MSE that remains sufficiently lower than the shuffled control (see the example in Figure 3e). Our approach thus considers them for TFBM integration, tolerating that their MSE increases as long as it remains reasonably lower than the simulated control.

We then focused on the sets of genes for which we specifically integrate TFBM in one approach but not the other, and computed precision and recall curves of the corresponding sub-networks. The results (Figures S10c and S10d) showed better performance of our scheme, which suggests that the sets of genes considered for data integration on the basis of a comparison to a null hypothesis are more relevant than simply minimizing the MSE. Concordant results were obtained when computing the global qualities of inferred GRN. We observed a slight median MSE increase in our approach as compared to the minimal MSE approach (Figure S10a), while precision and recall against DAP-Seq are both globally improved in the two models by our integration scheme, especially recall (Figure S10b).

### DIOgene improves the modelling of nitrate signalling

Finally, we assessed the ability of the inferred GRN to model nitrate induction pathways in Arabidopsis roots by comparing them to state of the art knowledge about this well documented response [8, 60]. In order to identify the regulators predicted as important players in nitrate response by our models, we ranked regulators by out-degree in the inferred GRN. This was done for both weightedLASSO and weightedRF, either in GRN inferred with a global value of *α* = 0, *α* = 1, or with the proposed gene-specific optimisation of *α* (Figure S11).

A first observation is that, regardless of the chosen model or data integration strategy, the 25 TF with highest out-degree contain previously known master regulators of nitrate response. This includes DIV1 [15], TGA1 and TGA4 [2], as well as the homologs HHO2 and HHO3, belonging to the NIGT family and identified as repressing the expression of crucial nitrate transport genes [29, 50]. Interestingly, we also uncover VRN1 and CRF4 as connectivity hubs in all inferred GRN. These regulators were respectively proposed as candidate and validated actors in nitrate signalling pathways in the studies that generated the transcriptomic data used here [59, 11]. Overall, whole-GRN measures of gene connectivity showed that genes involved in the regulation of nitrate pathways, nitrate uptake, transport and metabolism (Table S3) have a significantly higher total degree than other genes, in both globally optimized (at *α* = 0 and *α* = 1) and gene-specifically optimized GRN (Figure S12).

On another hand, we noticed that gene-specific calibration of data integration uniquely retrieves important regulators of nitrate nutrition, that were not present in the 25 most connected TF of the inferred GRN with a global *α* (*α* = 0 or *α* = 1). In the case of weightedLASSO, only the proposed gene-specific data integration strategy retrieves NLP7, which has been intensively documented as one of the main orchestrator of the early nitrate response [40, 3]. This is also the case of PHL1, a TF involved in the links between nitrate and phosphate signalling via NIGT-mediated regulations [58]. In the case of weightedRF, the proposed gene-specific optimization of data integration puts forward new TF as interesting candidates for nitrate response regulation. This includes HHO6, a member of the NIGT family not yet characterized for its role in the response to nitrate [29, 50], but also BZIP53, a TF involved in the regulation of several facets of metabolism [20] and JKD, a TF involved in shoot and root morphogenesis [30]. Thus, this analysis reveals that this method of inference, via the optimization of data integration in a gene-specific manner, not only recovers the information previously returned in the literature, but also brings to light new factors likely to be involved in this response.

## Discussion

The helpfulness of data integration is very often taken as granted in systems biology. Our work shows that it can in fact have very diverse effects on the modelling of gene expression, and that TFBM incorporation can be at the expense of model predictive capabilities for a significant number of genes. We thus propose to replace bulk data integration by a finely tuned hypothesis-driven data integration, calibrated individually for each gene. Our optimisation scheme, DIOgene, leverages TFBM in a way that their joint use with gene expression improves the target gene expression prediction over a simulated null hypothesis. On a plant biology case study, GRN inferred with this approach retrieve more DAP-Seq interactions through the use of TFBM and relevant known nitrate players, while preserving a near-optimal predictive performance on gene expression. Moreover, such conclusions hold for both the linear and non-linear regression cases, showing some general applicability to the most common models in the field. We also outlined some specificities in the tuning of TFBM integration between weightedLASSO and weightedRF (Figures S6, S11). Exploring these differences and the structure of the corresponding GRN would be a great way to test the impact of linearity and parametric assumptions in the modelling of multi-omics GRN.

The reason why some genes do not benefit from TFBM integration could stem from various factors, either technical or biological. Mining the consensual lists of genes for which *α* = 0 or *α >* 0 in both models revealed that genes for which TFBM are not integrated have in average a lower expression variance and more TFBM in their promoter (Figure S13). Further work would be needed to formulate hypotheses about the underlying general regulatory mechanisms, and also to assess the role of other forms of regulations like post-transcriptional and post-translational modifications in these results.

Several limitations of this study should be reminded to the reader. First of all, as in all works inferring GRN from expression data alone, the expression of the regulators is taken as a proxy for their activity. This assumption is not always valid, which motivated the estimation of TF activities in other studies, typically leveraging motifs or binding experiments combined to gene expression [34, 5, 22]. Our form of data integration, where TFBM-supported regulators have a stronger contribution in the estimated model, is another way to move away from this limitation. Even though this is a step toward more causality, challenges remain. One of them is the lack of a significant number of PWM, a problem amplified in non-model organisms. This limitation should be further reduced as PWM databases are completed and maintained by the community in the years to come, or as new computational methods are developed to predict binding affinities directly from DNA and protein sequences [7]. In the meantime, almost two thirds of regulators are attributed a neutral prior value, 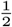in our study. The choice for this neutral prior could however be different and reflect other modelling strategies that we did not explore. Another limitation of the TFBM prior is the high chance of false positives when scanning PWM: the presence of a binding motif can occur by chance, or may not necessarily cause binding nor regulation in a cellular context. In addition, TFBM with low complexity in PWM databases can result in hits in almost all the promoters of an organism (Figure S4b). In this analysis, regulators with such a widespread PWM were included, but their questionable biological relevance could lead to their exclusion or to additional weighting based on PWM complexity during GRN inference. Moreover, the PWM themselves may not be accurate priors for TF binding, due to experimental and technical biases in their identification. Finally, non canonical binding events driven by features like DNA shape, structure or repeat sequences without the need for a motif have been reported [51], but cannot be directly modelled in our approach. Another limitation is the computation time of the proposed optimization procedure. Because it requires the estimation of a large number of models over a complete range of *α* values, computation time could be reduced by several ways such as a re-implementation in C++, or through analyses estimating the minimal number of repetitions to properly assess the MSE and EDI curves (Equations 6, 7, 9). Finally, strong levels of correlation in the input data are still hindering accurate GRN inference. A lot of pairs of regulators have correlated expression profiles. When their TFBM are different, our integration scheme can identify the relevant regulator; however when their TFBM are very similar or both unknown, identifying the meaningful one is not guaranteed. Correlation between variables also impacts the design of simulated null datasets: in our study we randomly unmatched the expression profiles of the regulators. This has the advantage of preserving the correlation structure in the expression data, thus creating a realistic null dataset. However, when many regulators have similar expression profiles, which is, to some extent, true in our case study, the simulated null data may sometimes partly resemble the original data only by chance. Bringing more diverse expression profiles into the simulated datasets, or constraining it so that it does not contain similar versions of the true data could be envisioned.

In addition to the aforementioned perspectives, the application of the proposed data integration strategy to other complex organisms is a promising lead. In this work, TFBM influencing gene expression were assumed to be located in the promoter regions of their target genes because very few distal regulations have been reported in Arabidopsis, and are still poorly understood [35]. In organisms where regulation by distant enhancers is well documented and responsible for tissue-specificity [4, 28], delineating enhancer regions may be achieved through the use of additional molecular layers such as chromatin accessibility, chromatin contacts, or eQTLs. Enhancers and promoters could then be scanned for TFBM, linked to their target genes and further guide GRN inference.

In our case study, we favored the use of model prediction performance as a quality metric because it is a condition specific metric available for all genes and orthogonal to the integrated TFBM priors, which is often not the case of current experimental gold standards. Our results indicate that instead of directly minimizing prediction error as a function of TFBM contribution, the comparison to a shuffled baseline improved inferred GRN (Figure S10). In essence, any inference method where data integration is tuned by a parameter could be optimized based on such a simulated null dataset. As a general guideline, we believe that both the monitored quality metric and the simulated baseline should be carefully designed in order to test a clear and relevant hypothesis for the problem at hand. More generally, the concept of synthetic null datasets for *in silico* negative controls is gaining interest in genomic analyses. For example, scDEED [61] optimizes 2D single-cell embeddings by simulating a null data where cells similarities are broken via permutations. Similarly, clusterDE [54] reduces false discoveries in marker gene identification by generating a realistic null single-cell data via scDesign3 [55], and then contrasting pipeline outputs between true and null datasets. Such synthetic controls, bearing similarities with the methodology proposed in this article, are likely to enhance rigor and causal discoveries in the field.

## Data and code availability

Below are the links to the data used in the course of this study.

- The RNA-Seq data for the response to nitrate induction was downloaded from the GEO accession GSE97500
- The PWM used to build the TFBM dataset were retrieved from JASPAR and the Plant Cistrome Database.
- To identify Arabidopsis TSSs and promoter regions, we relied on the TAIR10 GFF3 file.
- The regulators of Arabidopsis used for GRN inference are the union between PlnTFDB and AtTFDB

All results can be reproduced with the code available in the github repository: https://github.com/OceaneCsn/integrative_GRN_N_induction

## Author contributions statement

SL, LB, CHL and OC conceived the statistical inference methods and optimization scheme. OC implemented the inference methods and the optimization scheme. AM, SL and OC compared the inferred GRN to state of the art knowledge about nitrate signalling. All authors read and took part in writing the manuscript.

## Acknowledgments

This research is supported by a CNRS 80 Prime grant (BREAK), the French National Research Agency, and by the LabMUSE EpiGenMed (R-loops project). We thank Anaïs Baudot for her comment on an oral presentation of this work, which inspired a part of the discussion. We are grateful to all members of the Computational Regulatory Genomics team, Montpellier, for discussions around this project.

## Supplementary figures

**Table S1** : Normalized gene expression of the 1426 nitrate-responsive genes in the different treatments and time points of the experiment [59]. C: control. N: nitrate induction treatment. Numbers represent time after treatment in minutes.

**Table S2** : Gene identifier (AGI) of the 201 nitrate-responsive regulators.

**Table S3** : 56 genes involved in the uptake, transport, metabolism and signalling (positive or negative) of nitrate in the roots of Arabidopsis thaliana. Their AGI, gene name and short description are shown. This list of nitrate-related genes was compiled from the literature [8, 60, 15, 2, 29, 50, 59, 11, 40, 3, 58, 50, 20, 30].

**Fig. S1:**
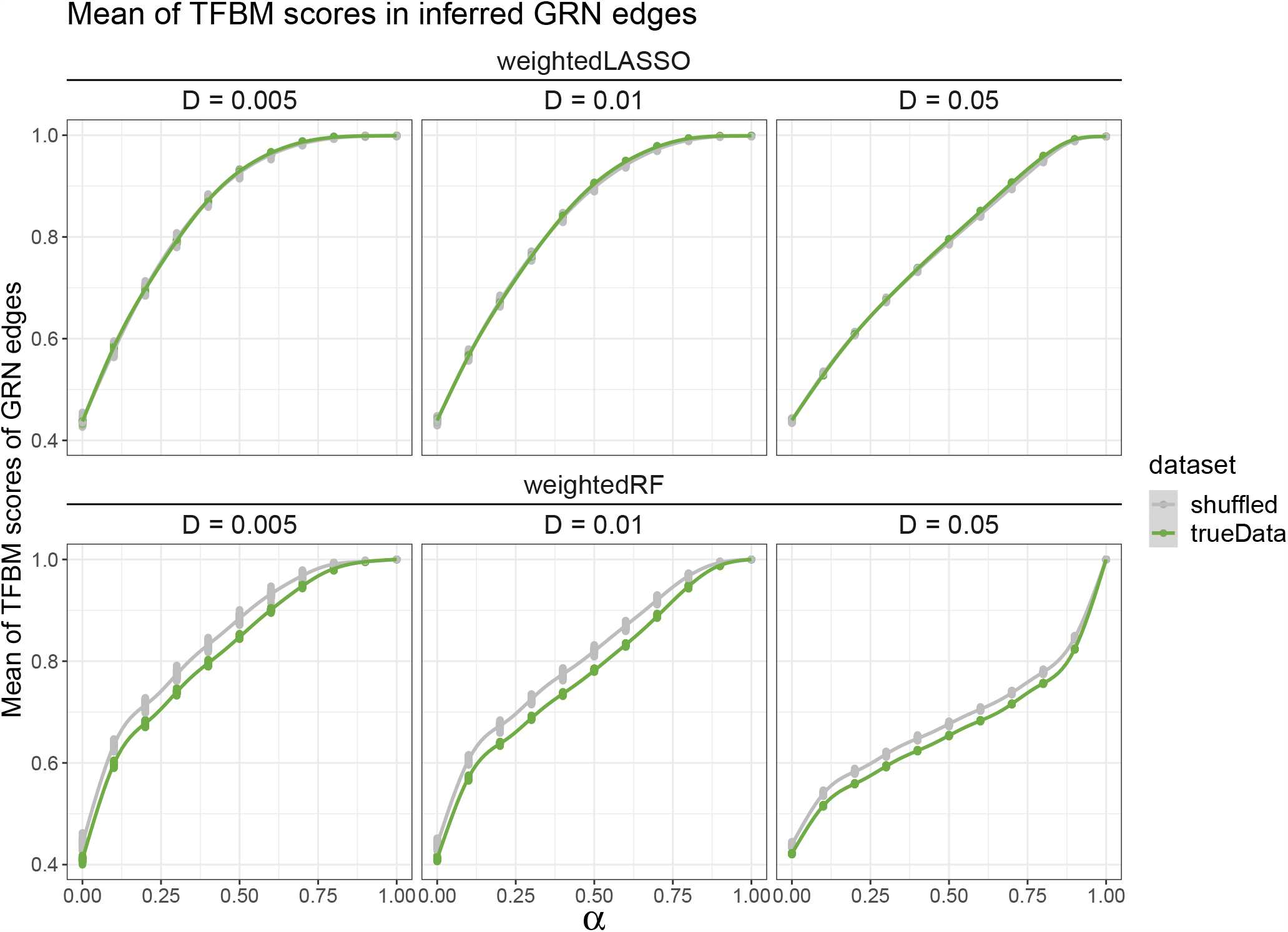
TFBM support of inferred GRN. TFBM support is the average value of the inferred TF-target edges in the prior TFBM matrix Π. It is shown for three network densities : 0.005 (1432 edges), 0.01 (2864 edges) and 0.05 (14322 edges). A TFBM of 1 means that a GRN is composed only of edges supported by a TFBM. At maximal TFBM integration intensity (*α* = 1), GRN on both true and shuffled data achieve a TFBM support of 1.

**Fig. S2:**
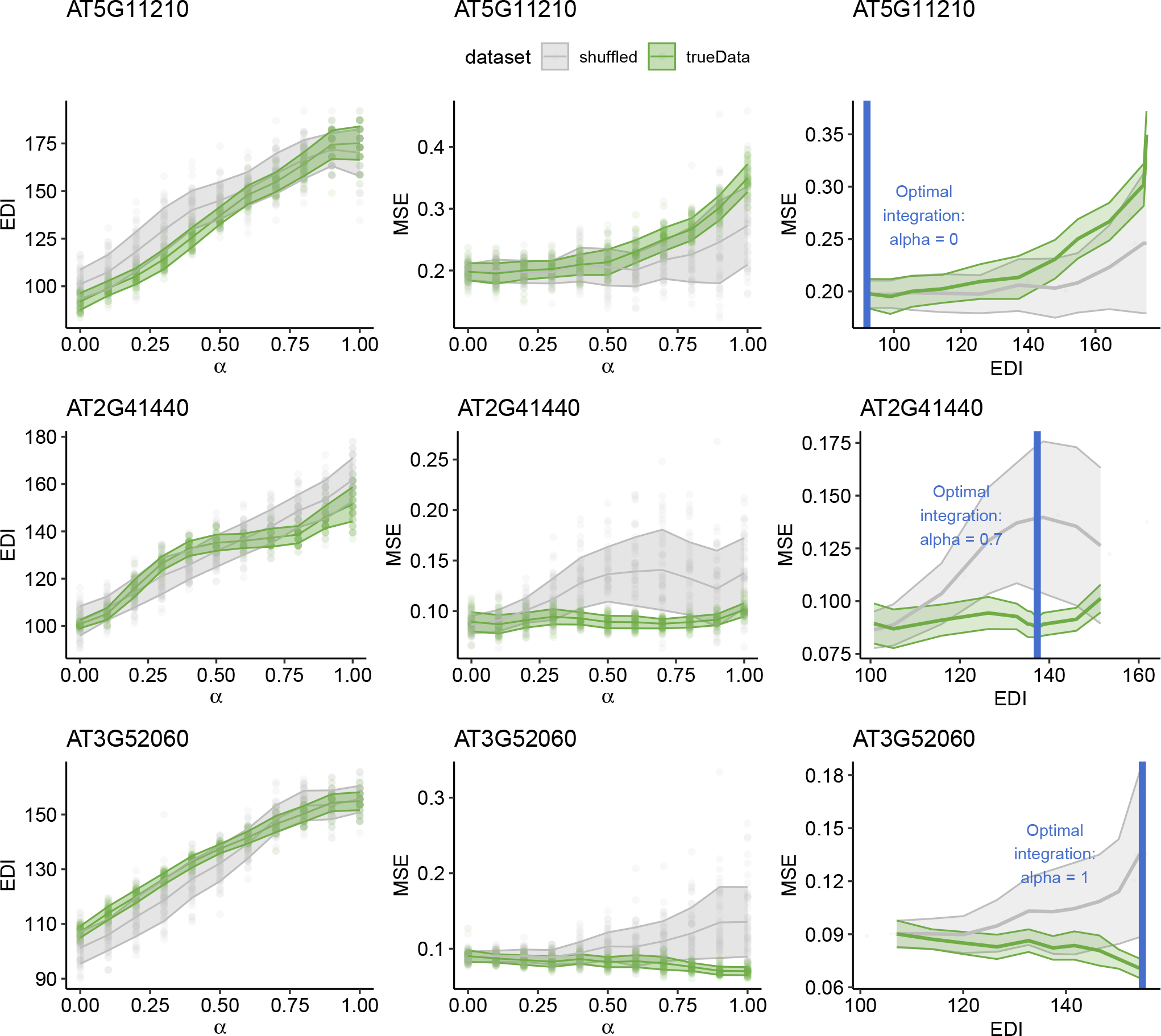
Different scenarios of data integration in weightedLASSO. For three gene examples in rows, the subplots show the EDI depending on *α*, the MSE depending on *α* and MSE depending on EDI for all possible values of *α* on true data (green) and shuffled null datasets (grey). The MSE is normalized by the variance of the target gene expression. For each value of *α*, 50 models were estimated and one standard deviation around the mean is represented. The proposed gene-specific *α*_opt_ is the value for which the MSE is most reduced as compared to the shuffled baseline, represented as a vertical blue line.

**Fig. S3:**
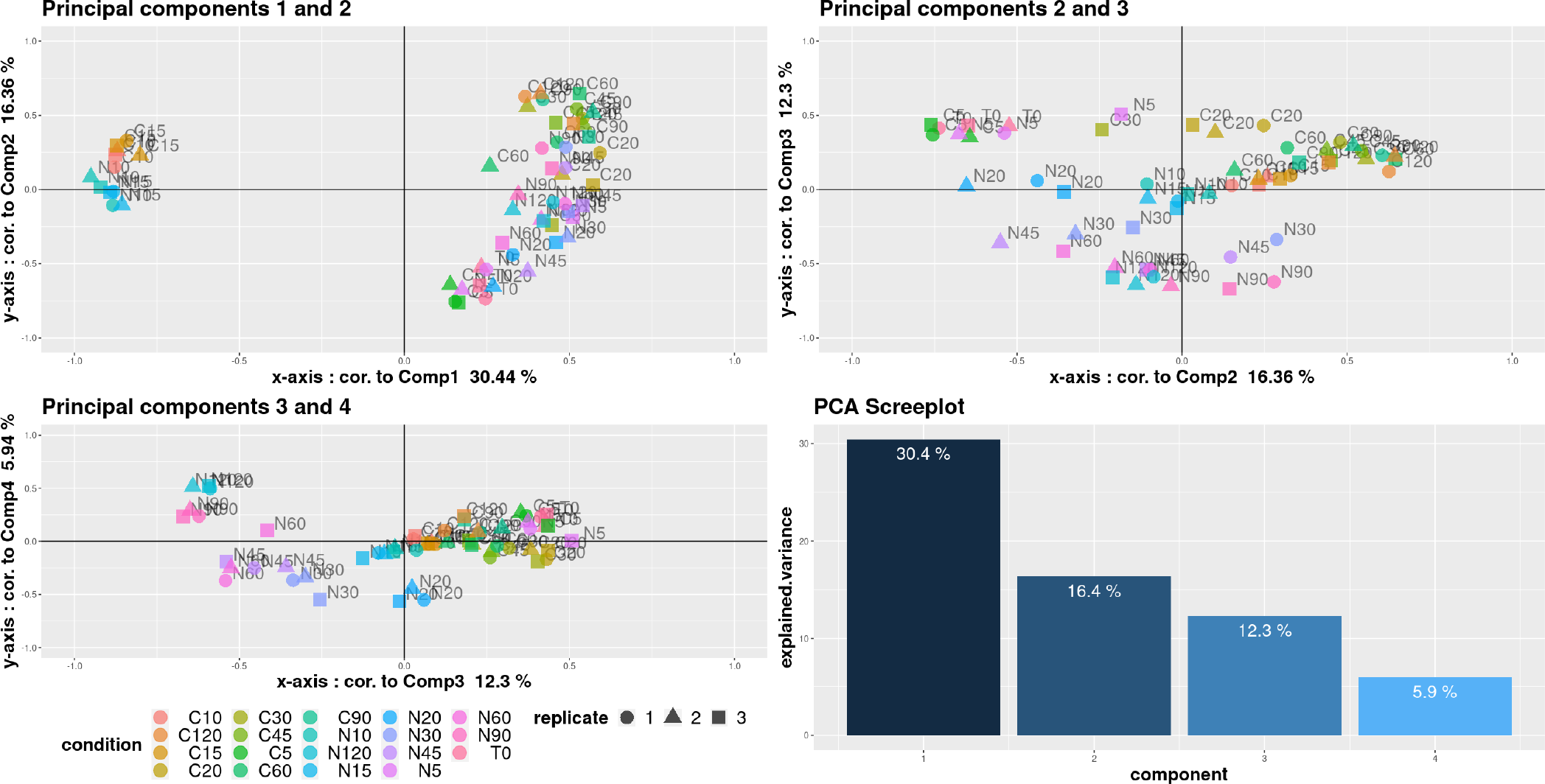
PCA plot of the normalized counts from the RNA-Seq experiment of the dynamic root response to a nitrate treatment [59]. The three first subplots show the correlation of each sample to the first 4 principal components. Each color represents an experimental condition made of 3 replicates. C: control. N: nitrate treatment. Numbers represent time in minutes. The first component highlights the samples at 10 and 15 minutes as different from the rest of the experiment, possibly due to a batch effect. The screeplot represents the percentage of variance explained by the 4 first principal components.

**Fig. S4:**
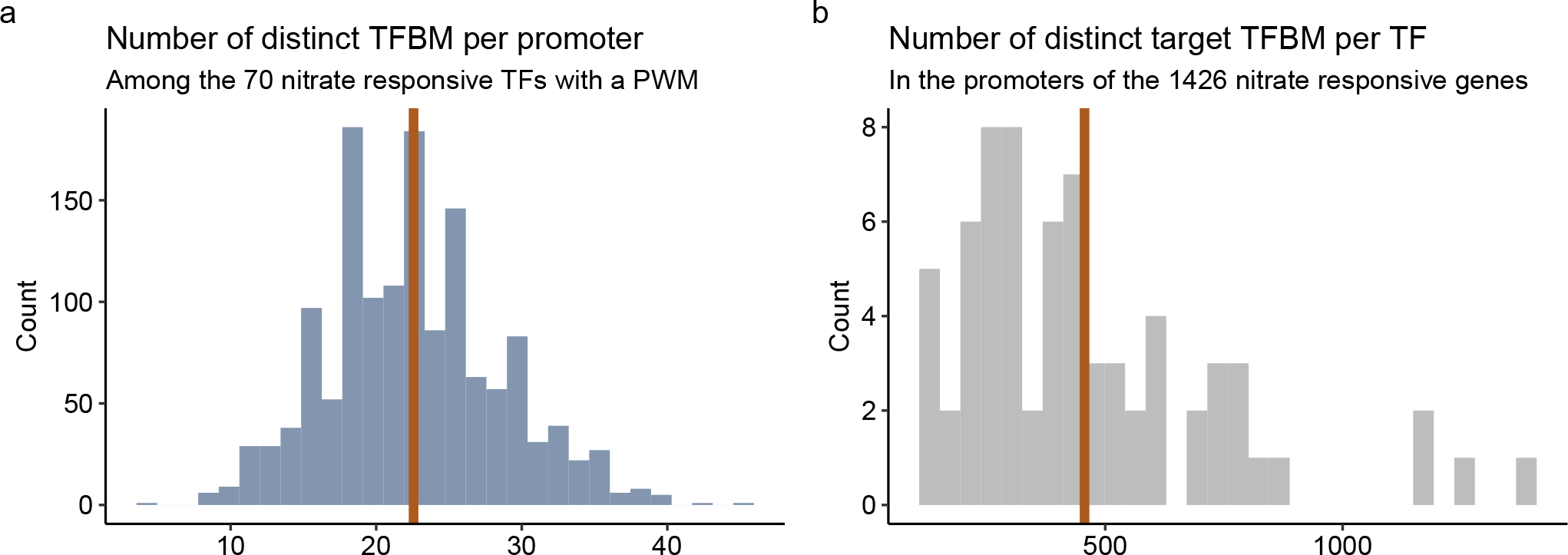
Prior TFBM network of nitrate-responsive genes (1426 promoters and 70 regulators). **a**. Distributions of the number of distinct PWM hits per promoter. **b**. Distribution of the number of distinct promoter hits per PWM restricted to nitrate-responsive genes. The average value of each distribution is represented by the orange vertical line.

**Fig. S5:**
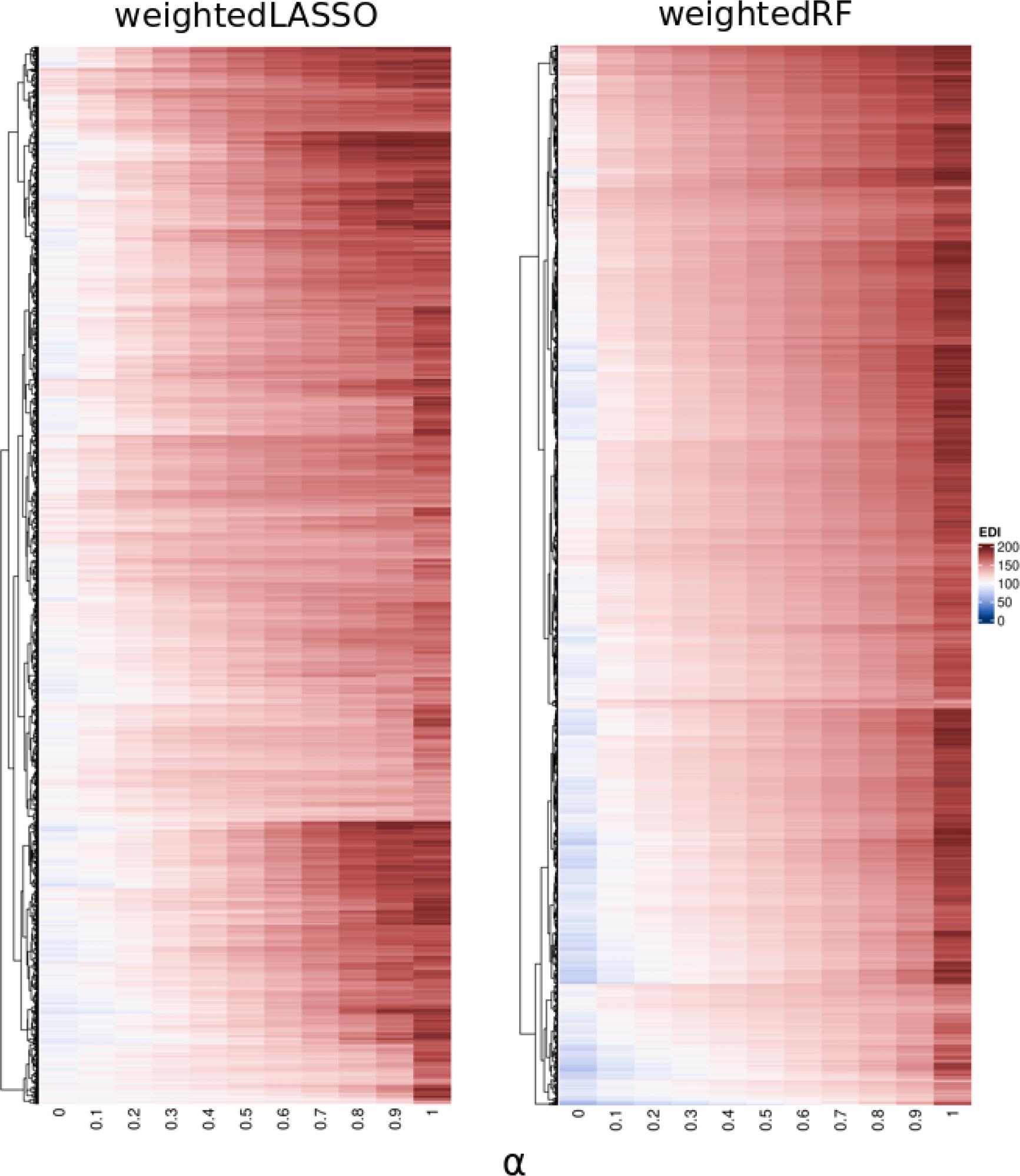
EDI for the 1426 nitrate-responsive genes as a function of *α*. Values are averaged across 100 replicates for weightedRF and 50

**Fig. S6:**
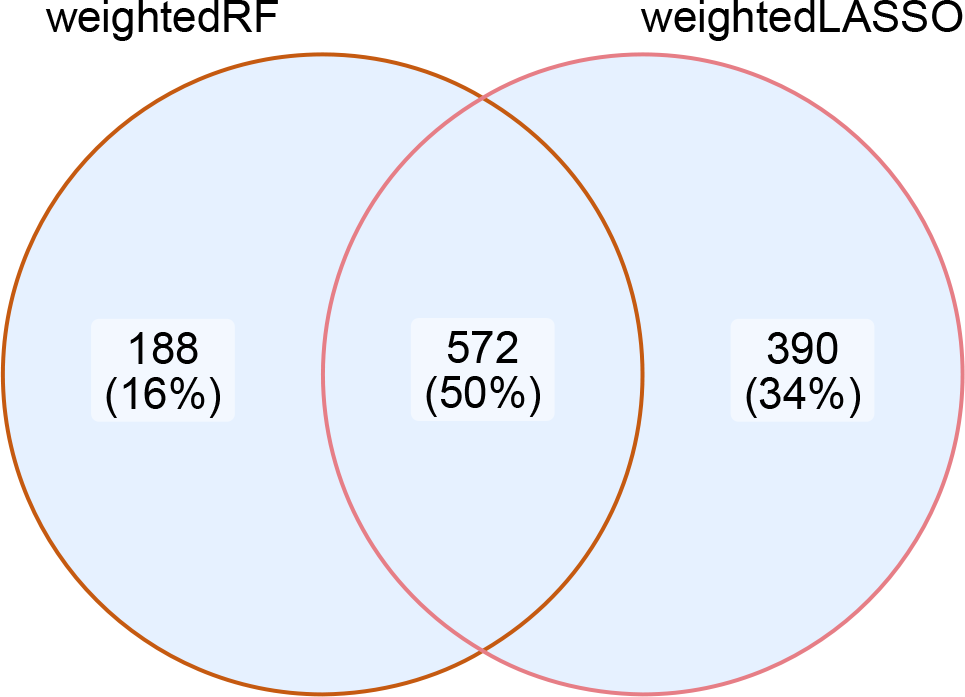
Intersection between the genes with *α*_*t,opt*_ *>* 0 in weightedLASSO and weightedRF with DIOgene. Enrichment pvalue : 5.9*e*^−12^ (One-sided hypergoemetric test).

**Fig. S7:**
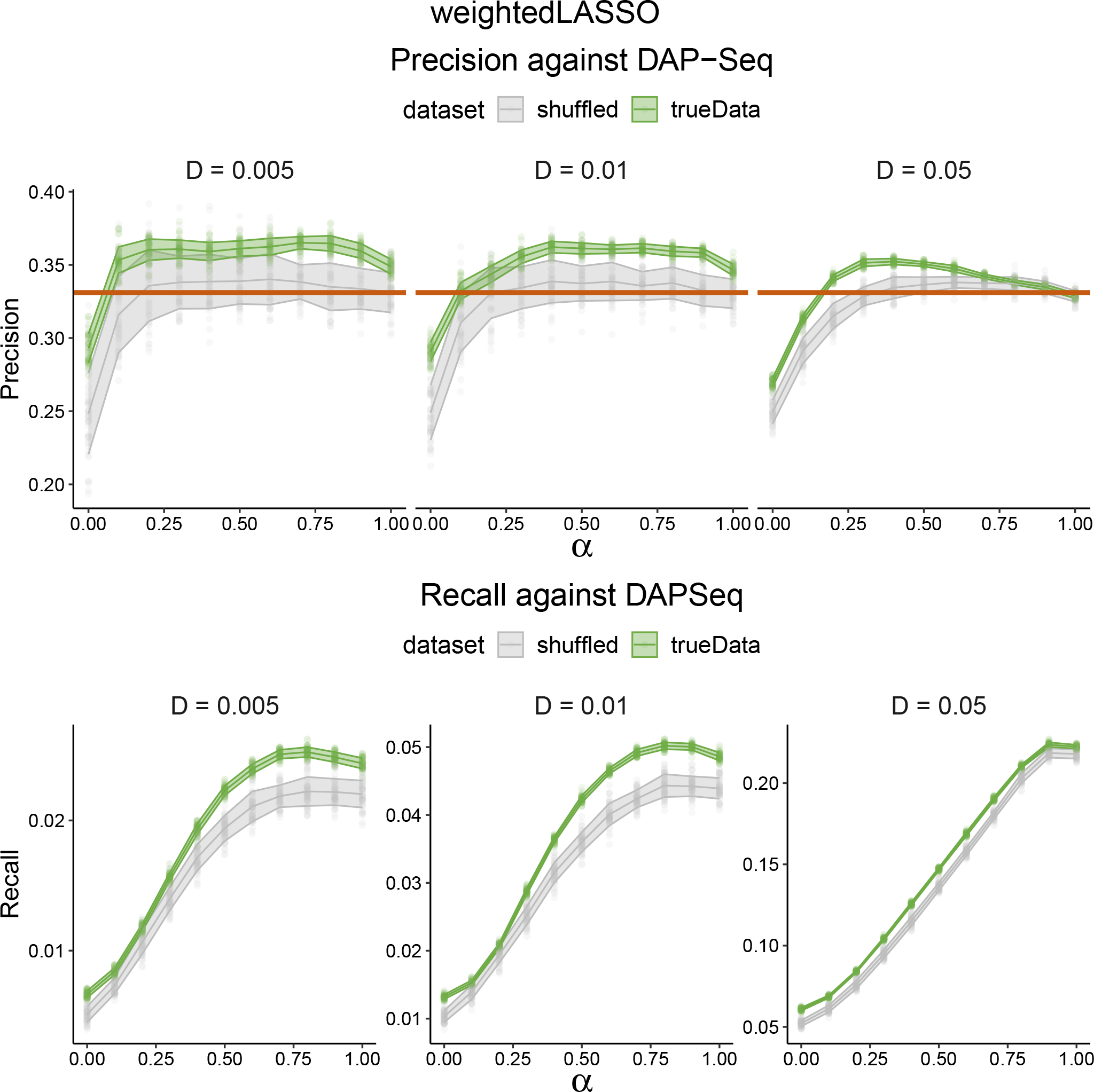
Precision and recall as a function of *α* in weightedLASSO, against a DAP-Seq gold standard. Precision and recall are show on true data (green) and shuffled null datasets (grey) for three network densities : 0.005 (1432 edges), 0.01 (2864 edges) and 0.05 (14322 edges). The precision of the prior TFBM network of nitrate responsive genes (31956 edges, density = 0.32) is overlaid in orange.

**Fig. S8:**
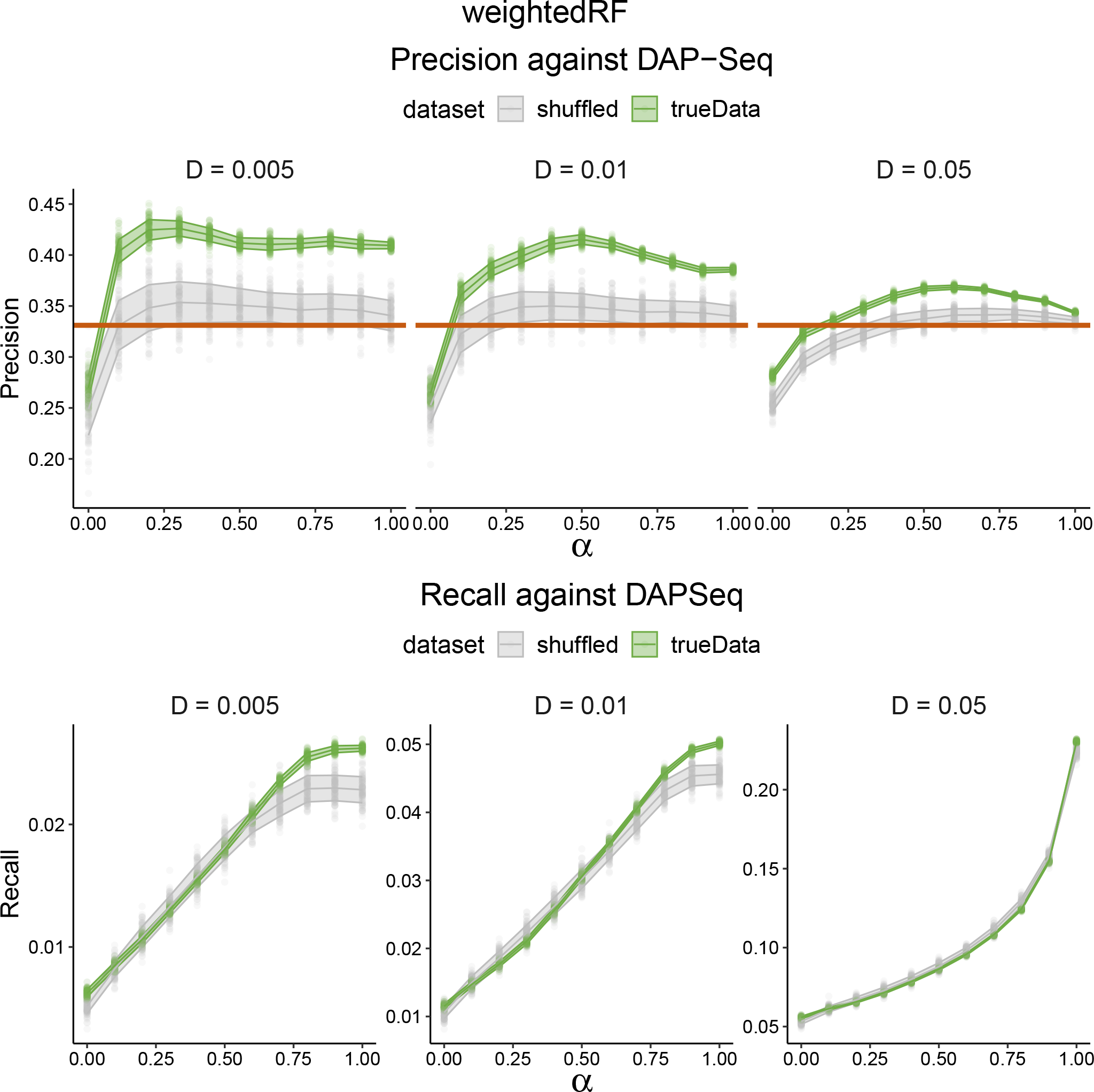
Precision and recall as a function of *α* in weightedRF, against a DAP-Seq gold standard. Precision and recall are show on true data (green) and shuffled null datasets (grey) for three network densities : 0.005 (1432 edges), 0.01 (2864 edges) and 0.05 (14322 edges). The precision of the prior TFBM network of nitrate responsive genes (31956 edges, density = 0.32) is overlaid in orange.

**Fig. S9:**
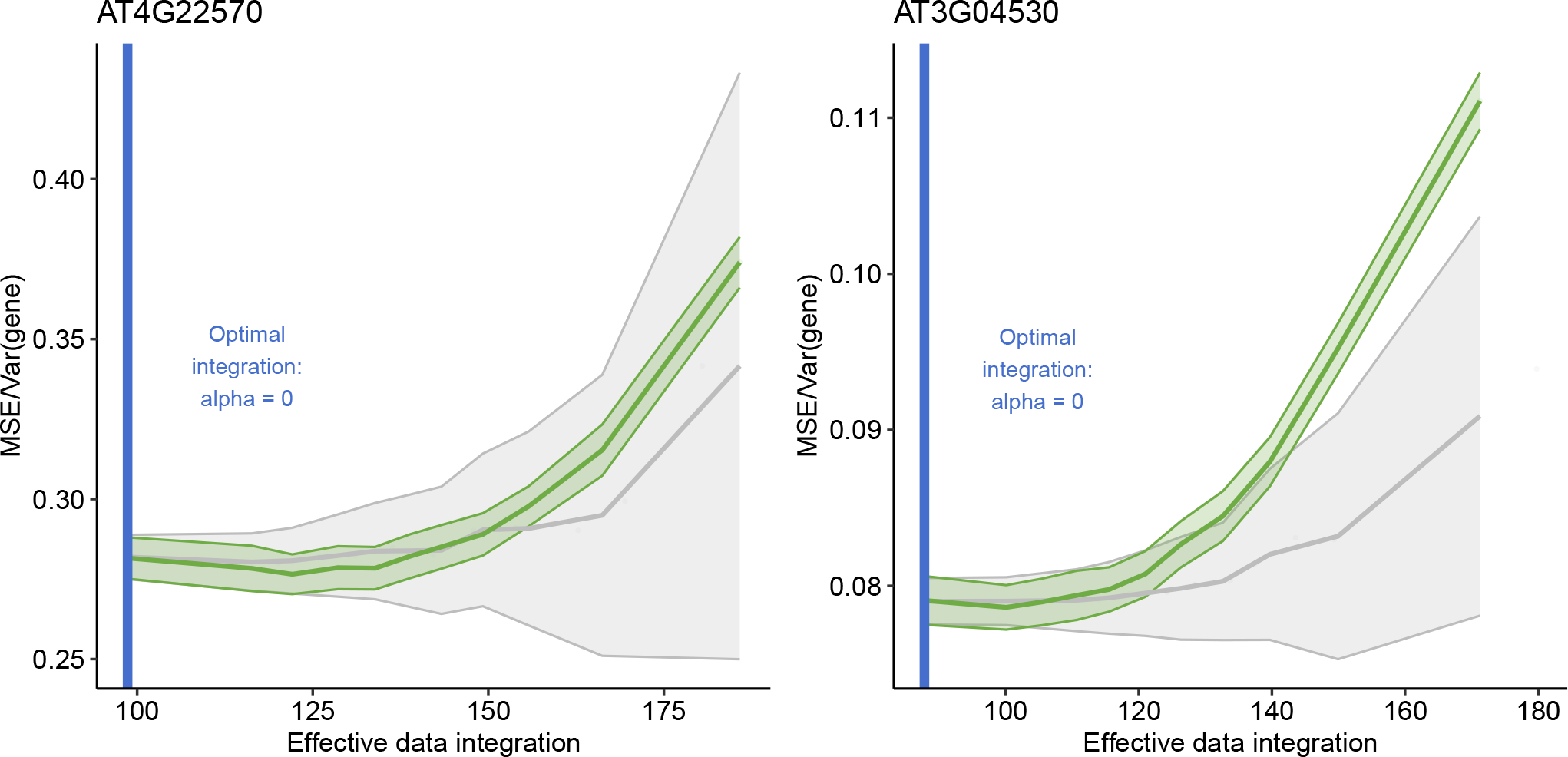
Example of two genes in weightedRF for which DIOgene sets *α*_*t,opt*_ = 0, but the minimal MSE approach would set *α*_*t,opt*_ *>* 0. MSE depending on EDI for all possible values of *α* on true data (green) and shuffled null datasets (grey).The MSE is normalized by the variance of the target gene expression.

**Fig. S10:**
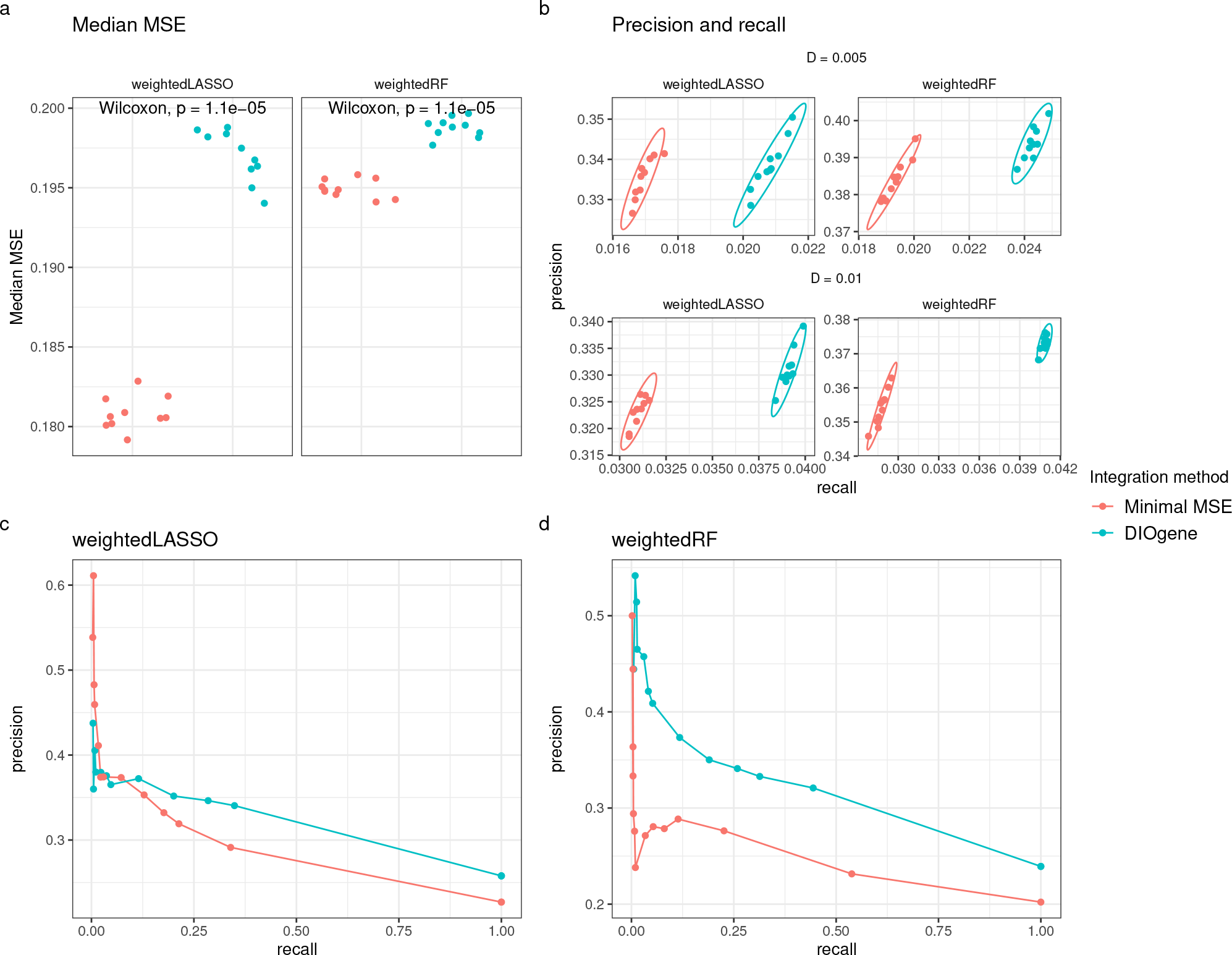
Comparison of the proposed gene-specific data integration tuning approach, DIOgene (blue), to the simpler minimal MSE approach (red). **a**. Median MSE, computed over all nitrate-responsive genes. The MSE is normalized by the variance of the target gene expression. **b**. Precision and recall computed against DAP-Seq interactions, on two GRN densities : 0.005 (1432 edges), 0.01 (2864 edges). Globally, the proposed tuning approach provides GRN closer to the DAP-Seq network. Each dot is a replicate of GRN inference, randomness stemming from different bootstrap sampling. **c-d**. Precision and recall curves for densities ranging from 0.001 to 1, computed on sub-GRN made of the edges concerning only target genes for which we integrate TFBM exclusively in one of the compared methods. In red, the genes for which the minimal MSE sets *α*_*t,opt*_ *>* 0 but not the proposed approach. In blue, the genes for which our approach sets *α*_*t,opt*_ *>* 0 but not the minimal MSE approach. This highlights the precision and recall gain from changing the sets of genes for which TFBM are integrated between the two methods.

**Fig. S11:**
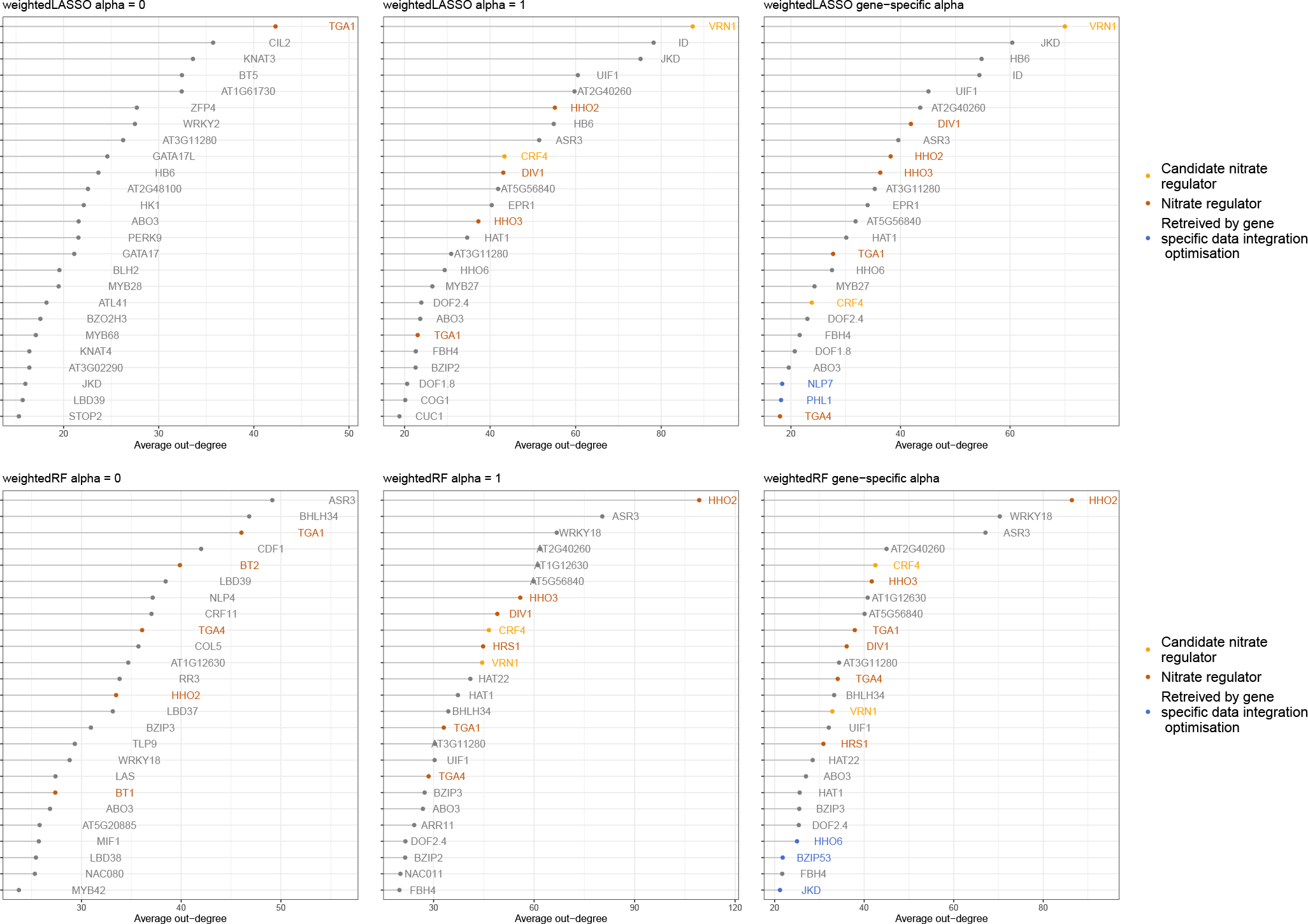
Most connected regulators depending on the model and TFBM integration strategy. In inferred GRN of a 0.005 density, regulators were ranked by their out-degree (number of regulated target genes or other regulators) for weightedLASSO and weightedRF, and for a global value of *α* (0 or 1), or using the proposed gene-specific approach (DIOgene). The out degree is averaged across the different runs of the same method to account for inherent stochasticity, and the top 25 TFs with the highest out-degree are shown. Regulators already identified in previous studies as important (orange) or candidate (yellow) actors of nitrate signalling and regulation are highlighted [8, 60, 15, 2, 29, 50, 59, 11, 40, 3, 58, 50, 20, 30]. TFs uniquely retrieved by the proposed gene-specific approach as compared to a global optimization for a given model are reported in blue.

**Fig. S12:**
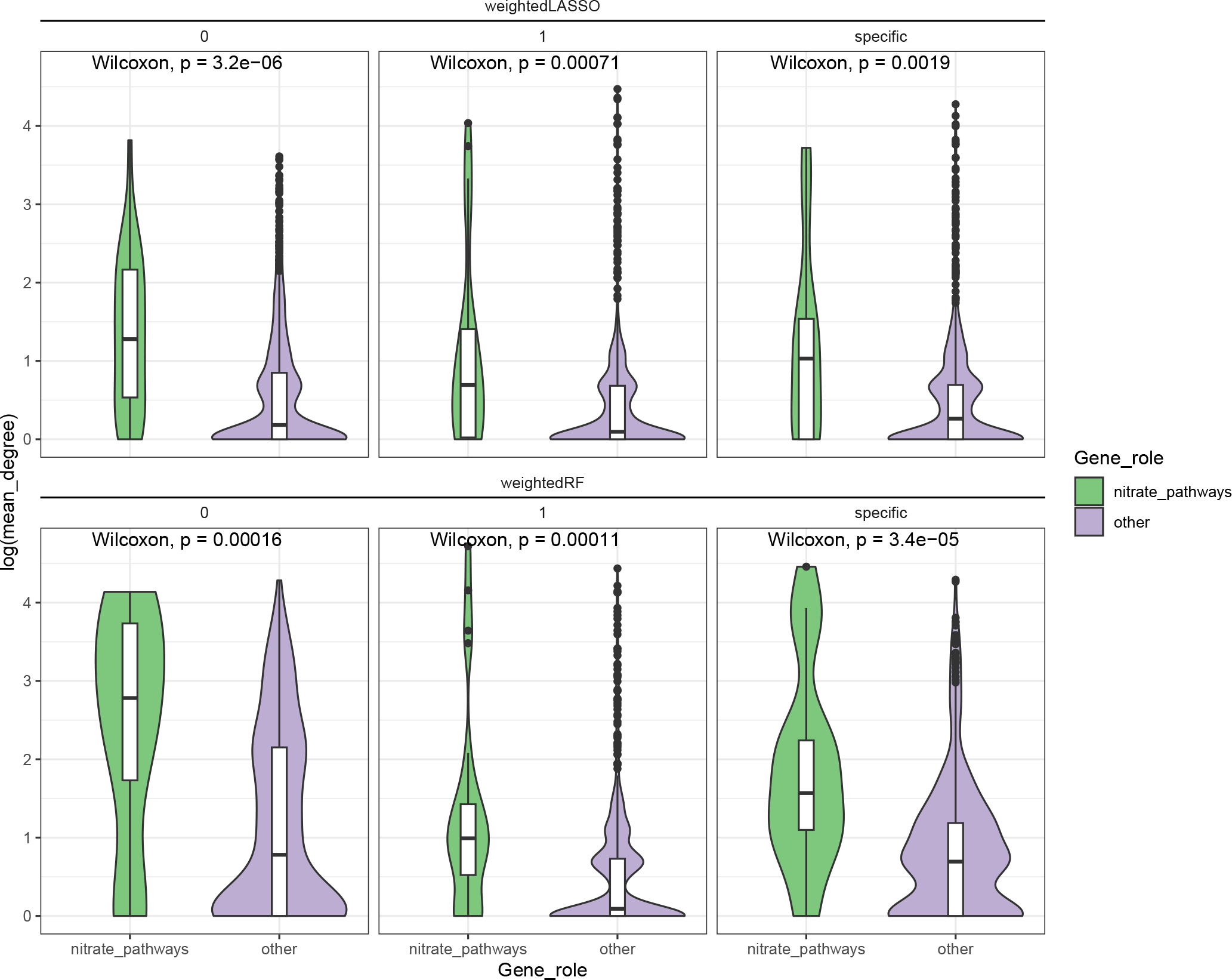
The overall degree (in-degree and out-degree) of nitrate-related genes is higher in inferred GRN as compared to other genes. In inferred GRN (*D* = 0.005), the total degree of genes is shown for a global value of *α* (0 or 1), or using DIOgene (specific). Total degree is reported on the log scale. Only genes with at least one connection in the inferred GRN were considered. The list of nitrate-related genes (Table S3) was compiled from the literature [8, 60, 15, 2, 29, 50, 59, 11, 40, 3, 58, 50, 20, 30].

**Fig. S13:**
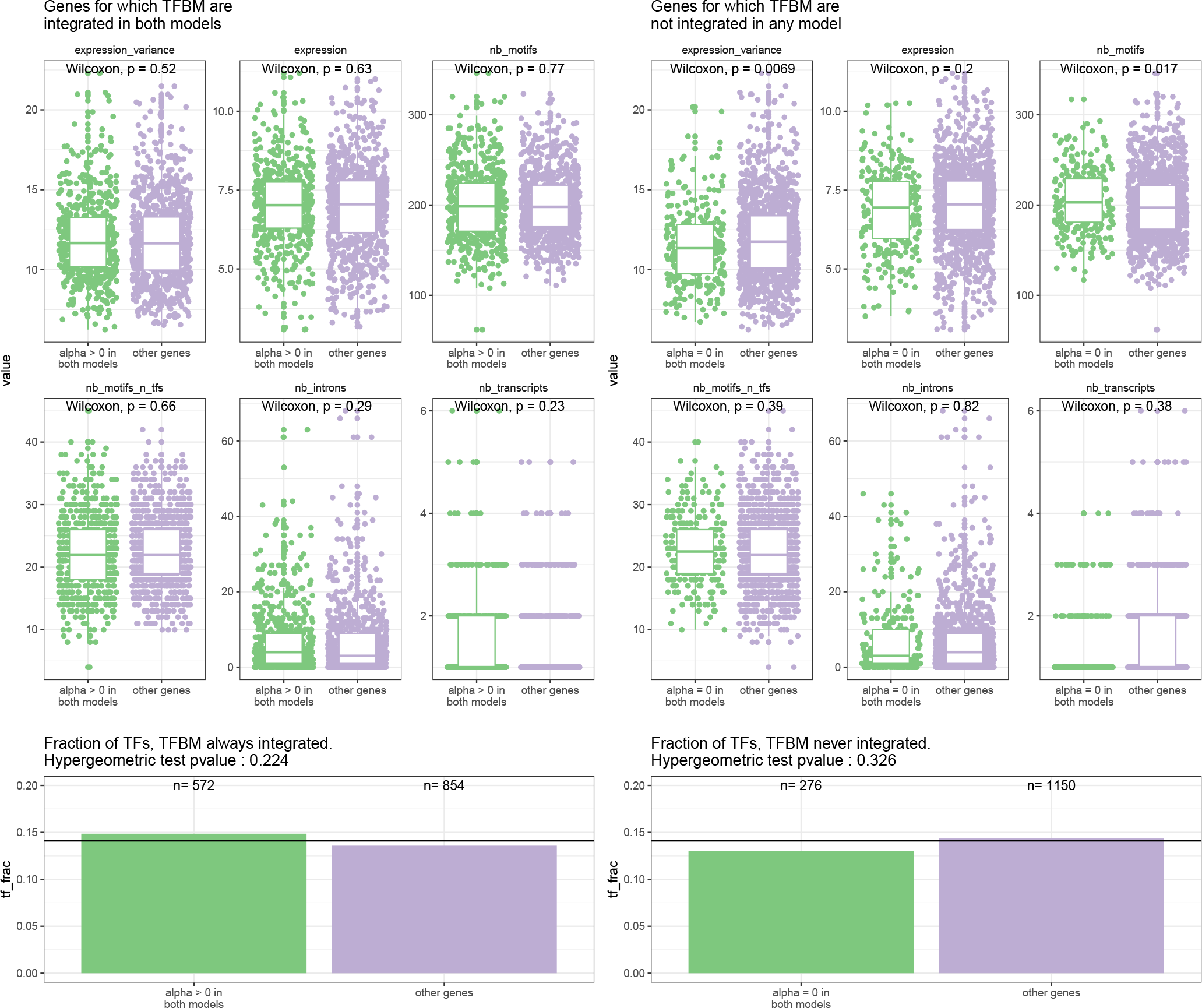
Functional comparison between genes for which TFBM are consensually integrated (*α*_*t,opt*_ *>* 0, left, *N* = 572) or not integrated (*α*_*t,opt*_ = 0, right, *N* = 276) in weightedRF and weightedLASSO. Expression and expression variance are reported on the log scale. The other characteristics are the number of motifs in the target gene promoter region, either among all knwon PWM in Arabidopsis (nb motifs), or only in nitrate responsive regulators (nb motifs n tfs), the number of introns (nb introns), the number of transcripts (nb transcripts). The two latter features were retrieved from the TAIR10 GFF annotation. On the last row, proportions of TFs in each group of genes are shown along an enrichment p-value result (one-sided hypergeometric test).

## Footnotes

1 Although samples at 10 and 15 min were also available, we discarded them, as they were clear outliers in the first dimension of a Principal Component Analysis (PCA) (Figure S3).

## References

1. Sara Aibar, Carmen Bravo González-Blas Thomas Moerman, Vân Anh Huynh-Thu, Hana Imrichova, Gert Hulselmans, Florian Rambow, Jean-Christophe Marine, Pierre Geurts, Jan Aerts, Joost van den Oord, Zeynep Kalender Atak, Jasper Wouters, and Stein Aerts. SCENIC: single-cell regulatory network inference and clustering. Nature Methods, 14(11):1083–1086, oct 2017.

2. José M. Alvarez Eleodoro Riveras, Elena A. Vidal, Diana E. Gras, Orlando Contreras-López, Karem P. Tamayo, Felipe Aceituno, Isabel Gómez, Sandrine Ruffel, Laurence Lejay, Xavier Jordana, and Rodrigo A. Gutiérrez. Systems approach identifies TGA1 and TGA4 transcription factors as important regulatory components of the nitrate response ofiarabidopsis thaliana/iroots. The Plant Journal, 80(1):1–13, aug 2014.

3. José M. Alvarez, Anna-Lena Schinke, Matthew D. Brooks Angelo Pasquino, Lauriebeth Leonelli, Kranthi Varala, Alaeddine Safi, Gabriel Krouk, Anne Krapp, and Gloria M. Coruzzi. Transient genome-wide interactions of the master transcription factor NLP7 initiate a rapid nitrogen-response cascade. Nature Communications, 11(1), March 2020.

4. Robin Andersson,, Claudia Gebhard, Irene Miguel-Escalada, Ilka Hoof, Jette Bornholdt, Mette Boyd, Yun Chen, Xiaobei Zhao, Christian Schmidl, Takahiro Suzuki, Evgenia Ntini, Erik Arner, Eivind Valen, Kang Li, Lucia Schwarzfischer, Dagmar Glatz, Johanna Raithel, Berit Lilje, Nicolas Rapin, Frederik Otzen Bagger, Mette Jørgensen, Peter Refsing Andersen, Nicolas Bertin, Owen Rackham, A. Maxwell Burroughs, J. Kenneth Baillie, Yuri Ishizu, Yuri Shimizu, Erina Furuhata, Shiori Maeda, Yutaka Negishi, Christopher J. Mungall, Terrence F. Meehan, Timo Lassmann, Masayoshi Itoh, Hideya Kawaji, Naoto Kondo, Jun Kawai, Andreas Lennartsson, Carsten O. Daub, Peter Heutink, David A. Hume, Torben Heick Jensen, Harukazu Suzuki, Yoshihide Hayashizaki, Ferenc Müller, Alistair R. R. Forrest, Piero Carninci, Michael Rehli, and Albin Sandelin. An atlas of active enhancers across human cell types and tissues. Nature, 507(7493):455–461, March 2014.

5. Mario L. Arrieta-Ortiz, Christoph Hafemeister, Ashley Rose Bate, Timothy Chu, Alex Greenfield, Bentley Shuster, Samantha N. Barry, Matthew Gallitto, Brian Liu, Thadeous Kacmarczyk, Francis Santoriello, Jie Chen, Christopher D. A. Rodrigues, Tsutomu Sato, David Z. Rudner, Adam Driks, Richard Bonneau, and Patrick Eichenberger. An experimentally supported model of the Bacillus subtilis global transcriptional regulatory network. Molecular Systems Biology, 11(11):839, November 2015.

6. Francis R Bach. Bolasso: model consistent lasso estimation through the bootstrap. In Proceedings of the 25th international conference on Machine learning, pages 33–40, 2008.

7. Sandro Barissi, Alba Sala, Mi-losz Wieczór, Federica Battistini, and Modesto Orozco. DNAffinity: a machine-learning approach to predict DNA binding affinities of transcription factors. Nucleic Acids Research, 50(16):9105–9114, August 2022.

8. Fanny Bellegarde, Alain Gojon, and Antoine Martin. Signals and players in the transcriptional regulation of root responses by local and systemic n signaling in arabidopsis thaliana. Journal of Experimental Botany, 68(10):2553–2565, mar 2017.

9. Linn Cecilie Bergersen, Ingrid K. Glad, and Heidi Lyng. Weighted lasso with data integration. Statistical Applications in Genetics and Molecular Biology, 10(1), January 2011.

10. Richard Bonneau, David J. Reiss, Paul Shannon, Marc Facciotti, Leroy Hood, Nitin S. Baliga, and Vesteinn Thorsson. The Inferelator: an algorithm for learning parsimonious regulatory networks from systems-biology data sets de novo. Genome Biology, 7(5):R36, May 2006.

11. Matthew D. Brooks, Jacopo Cirrone, Angelo V. Pasquino, Jose M. Alvarez, Joseph Swift, Shipra Mittal, Che-Lun Juang, Kranthi Varala, Rodrigo A. Gutiérrez, Gabriel Krouk, Dennis Shasha, and Gloria M. Coruzzi. Network walking charts transcriptional dynamics of nitrogen signaling by integrating validated and predicted genome-wide interactions. Nature Communications, 10(1), apr 2019.

12. Adrian I. Campos and Julio A. Freyre-González. Evolutionary constraints on the complexity of genetic regulatory networks allow predictions of the total number of genetic interactions. Scientific Reports, 9(1), mar 2019.

13. Océane Cassan, Sophie Lebre, and Antoine Martin. Inferring and analyzing gene regulatory networks from multi-factorial expression data: a complete and interactive suite. BMC Genomics, 22(1), May 2021.

14. Jaime A Castro-Mondragon, Rafael Riudavets-Puig, Ieva Rauluseviciute, Roza Berhanu Lemma, Laura Turchi, Romain Blanc-Mathieu, Jeremy Lucas, Paul Boddie, Aziz Khan, Nicolás Manosalva Pérez, Oriol Fornes, Tiffany Y Leung, Alejandro Aguirre, Fayrouz Hammal, Daniel Schmelter, Damir Baranasic, Benoit Ballester, Albin Sandelin, Boris Lenhard, Klaas Vandepoele, Wyeth W Wasserman, François Parcy, and Anthony Mathelier. JASPAR 2022: the 9th release of the open-access database of transcription factor binding profiles. Nucleic Acids Research, 50(D1):D165–D173, nov 2021.

15. Chia-Yi Cheng, Ying Li, Kranthi Varala, Jessica Bubert, Ji Huang, Grace J. Kim, Justin Halim, Jennifer Arp, Hung-Jui S. Shih, Grace Levinson, Seo Hyun Park, Ha Young Cho, Stephen P. Moose, and Gloria M. Coruzzi. Evolutionarily informed machine learning enhances the power of predictive gene-to-phenotype relationships. Nature Communications, 12(1), September 2021.

16. Scott Christley, Qing Nie, and Xiaohui Xie. Incorporating existing network information into gene network inference. PLoS ONE, 4(8):e6799, aug 2009.

17. Jacopo Cirrone, Matthew D. Brooks, Richard Bonneau, Gloria M. Coruzzi, and Dennis E. Shasha. OutPredict: multiple datasets can improve prediction of expression and inference of causality. Scientific Reports, 10(1), April 2020.

18. Inge De Clercq, Jan Van de Velde, Xiaopeng Luo, Li Liu, Veronique Storme, Michiel Van Bel, Robin Pottie, Dries Vaneechoutte, Frank Van Breusegem, and Klaas Vandepoele. Integrative inference of transcriptional networks in arabidopsis yields novel ROS signalling regulators. Nature Plants, 7(4):500–513, apr 2021.

19. Jerome Friedman, Trevor Hastie, and Robert Tibshirani. Regularization paths for generalized linear models via coordinate descent. Journal of Statistical Software, 33(1):1–22, 2010.

20. Abhroop Garg, Tobias Kirchler, Sven Fillinger, Friederike Wanke, Bettina Stadelhofer, Mark Stahl, and Christina Chaban. Targeted manipulation of bZIP53 DNA-binding properties influences arabidopsis metabolism and growth. Journal of Experimental Botany, 70(20):5659–5671, June 2019.

21. Pierre Geurts et al. dyngenie3: dynamical genie3 for the inference of gene networks from time series expression data. Scientific reports, 8(1):1–12, 2018.

22. Claudia Skok Gibbs, Christopher A Jackson, Giuseppe-Antonio Saldi, Andreas Tjärnberg, Aashna Shah, Aaron Watters, Nicholas De Veaux, Konstantine Tchourine, Ren Yi, Tymor Hamamsy, Dayanne M Castro, Nicholas Carriero, Bram L Gorissen, David Gresham, Emily R Miraldi, and Richard Bonneau. High-performance single-cell gene regulatory network inference at scale: the inferelator 3.0. Bioinformatics, 38(9):2519–2528, feb 2022.

23. Charles E. Grant, Timothy L. Bailey, and William Stafford Noble. FIMO: scanning for occurrences of a given motif. Bioinformatics, 27(7):1017–1018, feb 2011.

24. Alex Greenfield, Christoph Hafemeister, and Richard Bonneau. Robust data-driven incorporation of prior knowledge into the inference of dynamic regulatory networks. Bioinformatics, 29(8):1060–1067, 2013.

25. Anne-Claire Haury, Fantine Mordelet, Paola Vera-Licona, and Jean-Philippe Vert. TIGRESS: Trustful inference of gene REgulation using stability selection. BMC Systems Biology, 6(1), nov 2012.

26. Wayne Hayes, Kai Sun, and Nataša Pržulj. Graphlet-based measures are suitable for biological network comparison. Bioinformatics, 29(4):483–491, 2013.

27. Vân Anh Huynh-Thu, Alexandre Irrthum, Louis Wehenkel, and Pierre Geurts. Inferring regulatory networks from expression data using tree-based methods. PloS one, 5(9):1–10, 2010.

28. Aryan Kamal, Christian Arnold, Annique Claringbould, Rim Moussa, Nila H Servaas, Maksim Kholmatov, Neha Daga, Daria Nogina, Sophia Mueller-Dott, Armando Reyes-Palomares, Giovanni Palla, Olga Sigalova, Daria Bunina, Caroline Pabst, and Judith B Zaugg. scpGRaNIE/scp and scpGRaNPA/scp: inference and evaluation of enhancer-mediated gene regulatory networks. Molecular Systems Biology, 19(6), April 2023.

29. Takatoshi Kiba, Jun Inaba, Toru Kudo, Nanae Ueda, Mineko Konishi, Nobutaka Mitsuda, Yuko Takiguchi, Youichi Kondou, Takeshi Yoshizumi, Masaru Ohme-Takagi, et al. Repression of nitrogen starvation responses by members of the arabidopsis garp-type transcription factor nigt1/hrs1 subfamily. The Plant Cell, 30(4):925–945, 2018.

30. Atsushi Kobayashi, Satoshi Miura, and Akiko Kozaki. INDETERMINATE DOMAIN PROTEIN binding sequences in the 5′-untranslated region and promoter of the SCARECROW gene play crucial and distinct roles in regulating SCARECROW expression in roots and leaves. Plant Molecular Biology, 94(1-2):1–13, March 2017.

31. Mikaela Koutrouli, Evangelos Karatzas, David Paez-Espino, and Georgios A. Pavlopoulos. A Guide to Conquer the Biological Network Era Using Graph Theory, jan 2020.

32. Kundaje, S. LIANOGLOU, X. Li, D. Quigley, M. Arias, C. H. Wiggins, L. Zhang, and C. Leslie. Learning regulatory programs that accurately predict differential expression with MEDUSA. Annals of the New York Academy of Sciences, 1115(1):178–202, oct 2007.

33. Robert D. Leclerc. Survival of the sparsest: Robust gene networks are parsimonious. Molecular Systems Biology, 4, 2008.

34. Yue Li, Minggao Liang, and Zhaolei Zhang. Regression analysis of combined gene expression regulation in acute myeloid leukemia. PLoS Computational Biology, 10(10):e1003908, October 2014.

35. Yuan Lin, Hainan Zhao, Magdalena Kotlarz, and Jiming Jiang. Enhancer-mediated reporter gene expression in iarabidopsis thaliana/i : a forward genetic screen. The Plant Journal, 106(3):661–671, March 2021.

36. Han Liu, Kathryn Roeder, and Larry Wasserman. Stability approach to regularization selection (stars) for high dimensional graphical models. Advances in neural information processing systems, 23, 2010.

37. Han Liu, Kathryn Roeder, and Larry Wasserman. Stability Approach to Regularization Selection (StARS) for High Dimensional Graphical Models. Advances in neural information processing systems, 24(2):1432–1440, December 2010.

38. Aviv Madar, Alex Greenfield, Harry Ostrer, Eric Vanden-Eijnden, and Richard Bonneau. The Inferelator 2.0: a scalable framework for reconstruction of dynamic regulatory network models. Annual International Conference of the IEEE Engineering in Medicine and Biology Society. IEEE Engineering in Medicine and Biology Society. Annual International Conference, 2009:5448–5451, 2009.

39. Daniel Marbach, Sushmita Roy, Ferhat Ay, Patrick E Meyer, Rogerio Candeias, Tamer Kahveci, Christopher A Bristow, and Manolis Kellis. Predictive regulatory models in drosophila melanogaster by integrative inference of transcriptional networks. Genome research, 22(7):1334–1349, 2012.

40. Chloé Marchive, François Roudier, Loren Castaings, Virginie Bréhaut, Eddy Blondet, Vincent Colot, Christian Meyer, and Anne Krapp. Nuclear retention of the transcription factor nlp7 orchestrates the early response to nitrate in plants. Nature communications, 4(1):1–9, 2013.

41. Nicolai Meinshausen and Peter Bühlmann. Stability selection. Journal of the Royal Statistical Society: Series B (Statistical Methodology), 72(4):417–473, 2010.

42. Emily R Miraldi, Maria Pokrovskii, Aaron Watters, Dayanne M Castro, Nicholas De Veaux, Jason A Hall, June-Yong Lee, Maria Ciofani, Aviv Madar, Nick Carriero, Dan R Littman, and Richard Bonneau. Leveraging chromatin accessibility for transcriptional regulatory network inference in T helper 17 cells. Genome Res., 29(3):449–463, March 2019.

43. K. K. Nicodemus and J. D. Malley. Predictor correlation impacts machine learning algorithms: implications for genomic studies. Bioinformatics, 25(15):1884–1890, may 2009.

44. Ronan C. O’Malley, Shao shan Carol Huang, Liang Song, Mathew G. Lewsey, Anna Bartlett, Joseph R. Nery, Mary Galli, Andrea Gallavotti, and Joseph R. Ecker. Cistrome and epicistrome features shape the regulatory DNA landscape. Cell, 165(5):1280–1292, may 2016.

45. Francesca Petralia, Pei Wang, Jialiang Yang, and Zhidong Tu. Integrative random forest for gene regulatory network inference. Bioinformatics, 31(12):i197–i205, jun 2015.

46. Jing Qin, Yaohua Hu, Feng Xu, Hari Krishna Yalamanchili, and Junwen Wang. Inferring gene regulatory networks by integrating ChIP-seq/chip and transcriptome data via LASSO-type regularization methods. Methods, 67(3):294–303, jun 2014.

47. Matthew E. Ritchie, Belinda Phipson, D. Wu, Yifang Hu, Charity W. Law, Wei Shi, and Gordon K. Smyth. limma powers differential expression analyses for RNA-sequencing and microarray studies. Nucleic Acids Research, 43(7):e47–e47, January 2015.

48. Mark D Robinson and Alicia Oshlack. A scaling normalization method for differential expression analysis of RNA-seq data. Genome Biology, 11(3):R25, 2010.

49. Sushmita Roy, Stephen Lagree, Zhonggang Hou, James A. Thomson, Ron Stewart, and Audrey P. Gasch. Integrated Module and Gene-Specific Regulatory Inference Implicates Upstream Signaling Networks. PLOS Computational Biology, 9(10):e1003252, October 2013. Publisher: Public Library of Science.

50. Alaeddine Safi, Anna Medici, Wojciech Szponarski, Florence Martin, Anne Clément-Vidal, Amy Marshall-Colon, Sandrine Ruffel, Frédéric Gaymard, Hatem Rouached, Julie Leclercq, Gloria Coruzzi, Benoît Lacombe, and Gabriel Krouk. GARP transcription factors repress arabidopsis nitrogen starvation response via ROS-dependent and -independent pathways. Journal of Experimental Botany, 72(10):3881–3901, mar 2021.

51. Md. Abul Hassan Samee. Noncanonical binding of transcription factors: time to revisit ispecificity/i? Molecular Biology of the Cell, 34(9), August 2023.

52. Erwan Scornet. Trees, forests, and impurity-based variable importance. arXiv preprint arXiv:2001.04295, 2020.

53. Alireza F. Siahpirani and Sushmita Roy. A prior-based integrative framework for functional transcriptional regulatory network inference. Nucleic Acids Research, 45(4):e21, February 2017.

54. Dongyuan Song, Kexin Li, Xinzhou Ge, and Jingyi Jessica Li. ClusterDE: a post-clustering differential expression (DE) method robust to false-positive inflation caused by double dipping. July 2023.

55. Dongyuan Song, Qingyang Wang, Guanao Yan, Tianyang Liu, Tianyi Sun, and Jingyi Jessica Li. scDesign3 generates realistic in silico data for multimodal single-cell and spatial omics. Nature Biotechnology, May 2023.

56. Matthew E. Studham, Andreas Tjärnberg, Torbjörn E.M. Nordling, Sven Nelander, and Erik L. L. Sonnhammer. Functional association networks as priors for gene regulatory network inference. Bioinformatics, 30(12):i130–i138, jun 2014.

57. Robert Tibshirani. Regression shrinkage and selection via the lasso. Journal of the Royal Statistical Society: Series B (Methodological), 58(1):267–288, 1996.

58. Yoshiaki Ueda, Takatoshi Kiba, and Shuichi Yanagisawa. Nitrate-inducible NIGT1 proteins modulate phosphate uptake and starvation signalling via transcriptional regulation of iSPX/i genes. The Plant Journal, 102(3):448–466, January 2020.

59. Kranthi Varala, Amy Marshall-Colón, Jacopo Cirrone, Matthew D. Brooks, Angelo V. Pasquino, Sophie Léran, Shipra Mittal, Tara M. Rock, Molly B. Edwards, Grace J. Kim, Sandrine Ruffel, W. Richard McCombie, Dennis Shasha, and Gloria M. Coruzzi. Temporal transcriptional logic of dynamic regulatory networks underlying nitrogen signaling and use in plants. Proceedings of the National Academy of Sciences, 115(25):6494–6499, may 2018.

60. Elena A. Vidal José M. Alvarez, Viviana Araus, Eleodoro Riveras, Matthew D. Brooks, Gabriel Krouk, Sandrine Ruffel, Laurence Lejay, Nigel M. Crawford, Gloria M. Coruzzi, and Rodrigo A. Gutiérrez. Nitrate in 2020: Thirty years from transport to signaling networks. The Plant Cell, 32(7):2094–2119, mar 2020.

61. Lucy Xia, Christy Lee, and Jingyi Jessica Li. scDEED: a statistical method for detecting dubious 2d single-cell embeddings. April 2023.

62. Chun-Ping Yu, Jinn-Jy Lin, and Wen-Hsiung Li. Positional distribution of transcription factor binding sites in arabidopsis thaliana. Scientific Reports, 6(1), apr 2016.

63. Hui Zou and Trevor Hastie. Regularization and variable selection via the elastic net. Journal of the Royal Statistical Society: Series B (Statistical Methodology), 67(2):301–320, 2005. _eprint: https://onlinelibrary.wiley.com/doi/pdf/10.1111/j.1467-9868.2005.00503.x.

